# Coordinated representational drift supports stable place coding in hippocampal CA1

**DOI:** 10.1101/2025.02.04.636428

**Authors:** Ole Christian Sylte, Antje Kilias, Marlene Bartos, Jonas-Frederic Sauer

**Affiliations:** Institute of Physiology 1, Medical Faculty, University of Freiburg; Faculty of Biology, University of Freiburg

## Abstract

The phenomenon of representational drift (i.e., changing neuronal tuning during repeated exposure to the same stimuli), is a fundamental paradox in neuroscience that raises the question how stable behaviour can emerge from unstable neural representations. Place cells in CA1 of the hippocampus are crucial for spatial navigation and memories^1^ but show gradual changes in their preferred firing location over time when exposed to the same environment^2–9^. This drift occurs despite the fact that the animal maintains the ability to navigate and to perform spatial tasks, suggesting a complex relationship between neural activity and behaviour. Here, we show that while the spatial tuning functions of individual CA1 neurons drift over time, stable space coding prevails at the population level: Representational drift is not random but can be described as a translation and rotation in population state space, which preserves the internal geometry of population activity over time. Compensating for the coordinated translation and rotation allows for drift correction and a recovery of spatial tuning on future days. Moreover, the preserved internal geometry stabilises downstream readout under noisy conditions. We propose that the conserved population geometry might serve as a mechanism by which downstream reader networks achieve effective drift correction and, thus, ameliorate the readout of stable information.

## Introduction

Representational drift describes dynamically changing neuronal tuning functions in relation to identical external stimuli. Varying levels of drift have been observed in sensory^10–13^, motor^14^ and associative neocortex^15–18^ as well as in different hippocampal subfields^3^. The hippocampal output region CA1, which contains a spatial map of the environment in the form of firing fields of place cells^19^, is also subject to drift. While CA1 place fields are stabilised by spatial learning^20,21^ and maintain consistent field locations in bats undergoing stereotypical flight paths^22^, the spatial map in this brain region shows prominent representational drift under a variety of behavioural conditions^2–9,23,24^. This drift effectively decorrelates the hippocampal map within days to weeks. Drifting spatial representations pose a major conceptual challenge: everyday experience lets us perceive the world in a continuous and stable way: Consider walking along a familiar street. This experience does not seem different from one week to the next. At the same time, the observable neural correlate (i.e., the collection of spatial tuning functions of hippocampal neurons) drifts over time and is reconfigured into a new spatial map.

How can downstream circuits extract stable spatial information from such drifting hippocampal maps? One solution to this problem might be that target networks are relatively unaffected by drift if it occurs in a ‘null’-space, meaning that the changing CA1 activity does not sufficiently alter the information ‘seen’ by the cells downstream. While this appealing mechanism has been suggested to play a major role in separating motor preparation from movement signals^25,26^, spatial map drift in the posterior parietal cortex (PPC) occurs in part outside of null-space dimensions^27^. Drift in null-space is therefore unlikely to be the sole mechanism by which stability can be transferred out from drifting circuits, and different communicating circuits might employ distinct methods of drift correction. Modelling work suggests that plasticity in the downstream network and comparisons to a reference signal might additionally alleviate the impact of drifting inputs^28^, but it is not known to what extent the networks downstream of CA1 implement such correction mechanisms. How drifting CA1 output might be converted to a temporally consistent signal, therefore, remains unclear.

Here, we explored a different way by which drifting output might be perceived as stable downstream of CA1: We propose that drift is not a random process (i.e., that the neurons’ tuning functions do not change independently for each cell) but that it occurs coordinated across individual neurons. This hypothesis states that a stable code for space is embedded in the *relationship* among neuronal firing patterns, rather than in the absolute firing properties and tuning functions of individual neurons. These relationships form an ‘internal geometry’ - a set of relative distances and angles between population responses that contains all decodable information. We show that this geometry remains stable over time, allowing downstream networks to read the same spatial map despite ongoing drift.

## Results

### Drifting CA1 place fields

To assess hippocampal place map stability, we performed multi-day 2-photon calcium imaging from CA1 pyramidal neurons of head-fixed mice exploring virtual linear tracks^3^ (**Fig. 1A,B**, Supplemental Fig. 1A-D). The animals were trained to collect liquid rewards at fixed track locations (Supplemental Fig. 1E-H). Due to conflicting reports on the magnitude of representational drift in CA1^20–22^, we first quantified the stability of the hippocampal map in our virtual navigation paradigm. Visualising the spatial tuning functions of all observed CA1 pyramidal neurons over 20 days on a familiar virtual track revealed a drifting place code (**Fig. 1C**). This was evident from a significant decrease in the correlation of spatial tuning functions over subsequent days (**Fig. 1D**), with less than 3% of neurons maintaining stable spatial tuning over days (across-days Pearson’s r ≥ 0.7, Supplemental Fig. 1I). Accordingly, when predicting the animal’s position on the track with decoding models trained on calcium transients from the first day, we found the decoding error to increase with the time between the training and the testing day and to quickly become indistinguishable from the spatial error obtained from shuffled control data (**Fig. 1E,F**). These results confirm previous accounts of unstable spatial population codes in CA1^2–9^.

**Fig 1.**
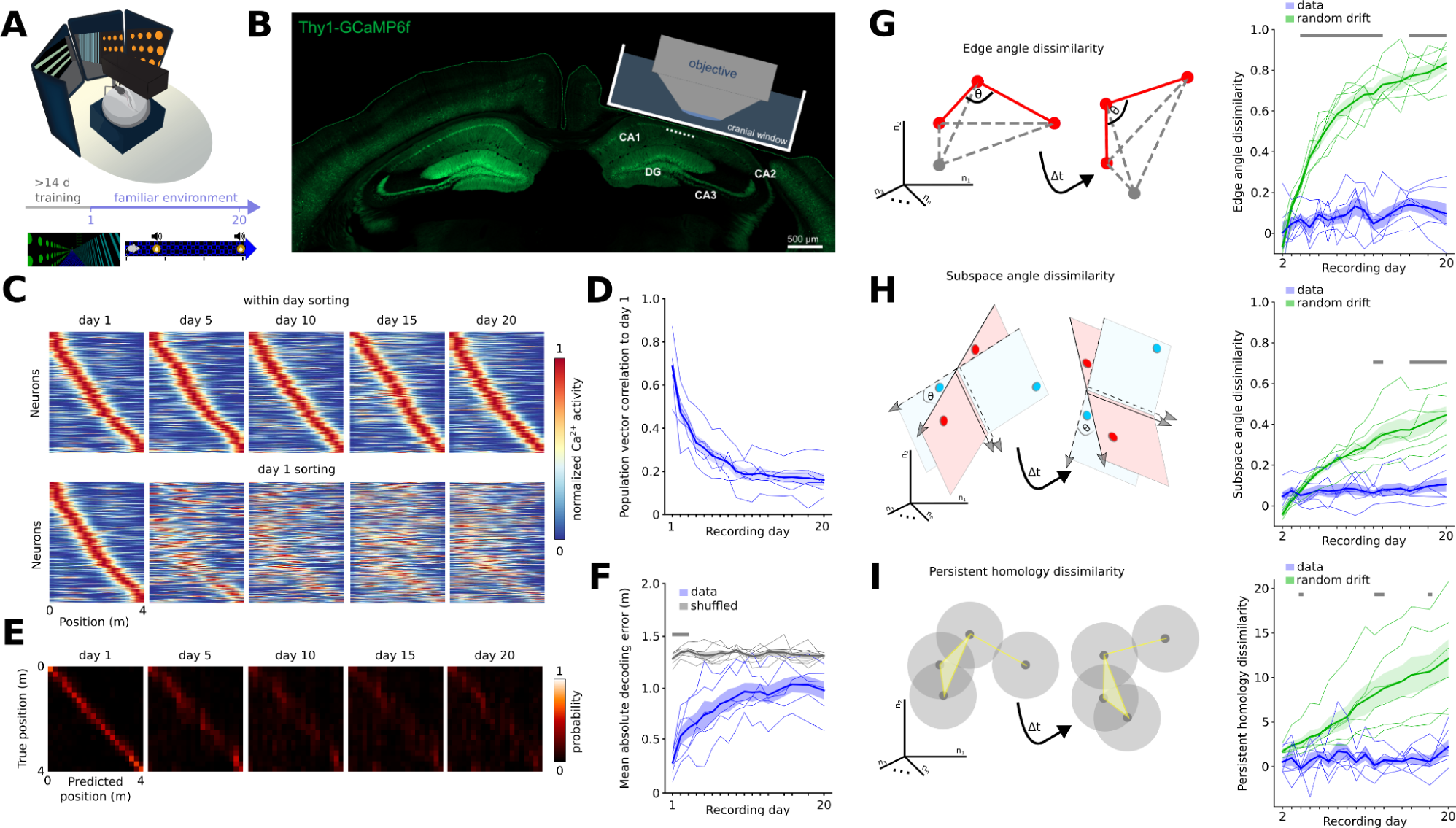
Stable internal geometry of population activity despite representational drift in CA1. **A.** Schematic of the recording configuration during virtual navigation along a linear track. **B.** Brain slice overlaid with a schematic cranial window implanted for 2P imaging from dorsal CA1 in Thy1-GCaMP6f mice. DG: dentate gyrus **C.** Spatial map (i.e., spatial tuning functions of CA1 pyramidal cells) of an example mouse sorted by peak activity within each day. Bottom: The same spatial map sorted by day 1. **D.** Population vector correlation to day 1 decreases over time. F=60.3, p=10^-38^, 1-way repeated measures ANOVA. Values at day 1 show within-day correlation. **E**. Gaussian Naive Bayes decoder trained on day 1 to predict the position of the animal based on calcium data of subsequent days. The data is averaged over all mice. **F**. Mean absolute error of the decoder vs. shuffled control data. t=6.3-15.1, p=0.026-10^-5^. **G.** Left: Illustration of edge angle (⊝) analysis to assess internal population geometry. Right: low edge angle dissimilarity persists relative to day 1. Data is scaled between 0 (within session stability) and 1 (complete shuffled data). Green: Data with simulated random drift. t=6.1-24.7, p=0.029-10^-5^. **H**. Same as G but for subspace angle analysis. t=5.4-7.6, p=0.042-0.010. **I.** Same as G but for persistent homology analysis. t=5.3-8.3, p=0.049-0.007. Grey bars in F, G, H, I: significant differences, 2-way repeated measures ANOVAs followed by paired *t* or Wilcoxon tests with Bonferroni correction. Thick lines/shading: mean ± SEM, thin lines: individual mice (n=6).

### Temporally stable internal geometry of population activity

Considering that the spatial map is subject to representational drift, we asked whether features of stability might be evident at the level of population activity. We hypothesised that the internal geometry of the population activity is preserved over time despite drift of the tuning functions of individual neurons. In other words, just as a constellation maintains its shape regardless of its orientation in the night sky, we propose that the drift of the individual cells’ activity in relation to space occurs in a coordinated rather than random fashion. For *N* recorded neurons, consider the geometrical object *G* defined by the activity of the population in *N-*dimensional space over *B* spatial bins. If the geometry of this object (i.e., the internal geometry of the population) is conserved over time, the angles between the connections among the *NxB* nodes will remain constant (**Fig. 1G**). In other words, a stable internal geometry might shift, rotate or uniformly stretch in its reference frame over subsequent days (i.e., undergo a similarity transformation), but it will remain the same geometrical object^10^. If, in contrast, drift occurs randomly for each neuron, the angles of *G* will change, and the cumulated difference in the measured angles will increase as a function of time, leading to a more dissimilar geometry. To take a reasonable lower bound of expected dissimilarity over days into account, we normalised *G* angle dissimilarity to the one observed between odd and even runs on the first recording day. We found that the edge angle dissimilarity of *G* constructed daily for a set of repeatedly active neurons on the familiar track remained low, even up to 20 recording days (**Fig. 1G**). To quantitatively judge the stability of *G*, we explored a reasonable upper bound of expected drift in case this drift occurred randomly rather than coordinated among the neurons: We performed the same analysis for a dataset in which *G* was constructed for neurons with simulated random drift in their place fields (Supplemental Fig. 2A). The time-dependent decorrelation of the drift simulation reflected the one observed for real data (Supplemental Fig. 2A). As expected, this model showed increasing edge angle dissimilarity over time, indicative of a cumulatively changing internal geometry (**Fig. 1G**). This effect was true for a range of simulated drift magnitudes (Supplemental Fig. 2B-D). Consistent internal geometries were preserved when we removed stable place cells from the population (Supplemental Fig. 2E), confirming that the stability of the internal geometry is not driven by the small proportion of temporally stable cells. To corroborate these findings, we used two complementary analyses to assess the internal geometry of the population: First, we examined the dissimilarity of local subspace angles across days (**Fig. 1H**). For this analysis, we extracted local subspaces for any two consecutive positions and computed the principal angles between these subspaces. As a second method, we used persistent homology to quantify abstract scale-invariant topological features (such as connected components, holes and voids) in the neural data that persist across various scales (**Fig. 1I**). We then compared the persistence diagram of these topological features across days. Both methods confirmed a stable population geometry across days compared to simulated random drift (**Fig. 1H,I**). Similar stabilities were observed when recomputing the geometry after projecting the data onto lower-dimensional spaces (Extended Fig. 2F,G). Taken together, CA1 population activity drifts in a coordinated, geometry-preserving manner.

**Fig 2.**
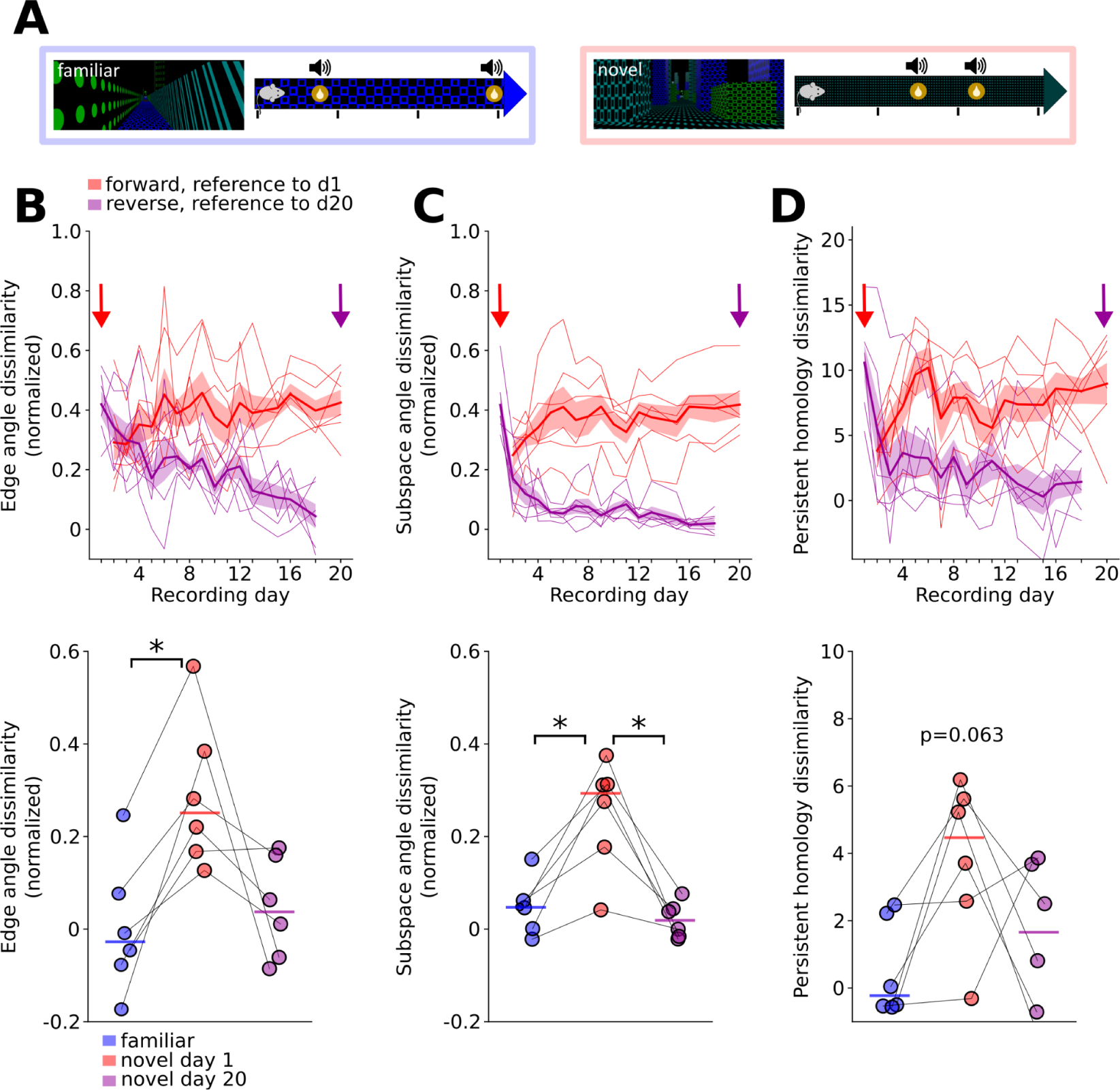
Experience-dependent stabilisation of internal geometry across days. **A**. Illustration of familiar and novel environments. **B**.Top: Edge angle dissimilarity in the novel environment relative to day 0 (‘forward’) and relative to day 20 (compared backwards in time, ‘reverse’). Data is scaled between 0 (within session stability) and 1 (complete shuffled data). Bottom: Across-day edge angle dissimilarity in the familiar environment (blue) and during the first (red) and last day of novel environment exposure (day 20, purple). Familiar vs. novel day 1: t=-4.8, p=0.015, all other comparisons p>0.05. **C.** Same as B but for subspace angle dissimilarity. Familiar vs. novel day 1: t=-4.6, p=0.017; familiar vs. novel day 20: t=1.9, p=0.346; novel day 1 vs. day 20: t=4.4, p=0.021. **D.** Same as B but for persistent homology dissimilarity. C,E,G: 1-way repeated measures ANOVA followed by paired *t* tests with Bonferroni correction. Thick lines/shading: mean ± SEM, thin lines: individual mice (n=6).

### Stable internal geometries emerge with experience

Previous work emphasised that the stability of the CA1 map increases with learning^21^. We therefore asked whether the stable internal geometry depends on prior experience with the track. To address this question, we explored how the geometric structure of the population evolves when the animals first encounter a novel environment. We recorded neural activity as mice explored a novel virtual track that differed from the familiar track in terms of visual features and reward locations (**Fig. 2A**). In contrast to the stable internal geometry on the familiar track, the internal population geometry in the novel environment underwent substantial changes during initial exposure, as indicated by larger edge angle dissimilarities relative to day 1 (**Fig. 2B**). In contrast, when we used the novel track on day 20 as a reference time point (i.e., when the animals were already familiar with this environment) and computed edge angle differences to previous days, we found low dissimilarity values instead (**Fig. 2B**). Similar results were obtained using the subspace angle dissimilarity, and we observed a trend towards the same outcome using the persistent homology dissimilarity measure (**Fig. 2C,D**). These data suggest that in a novel context, population geometry is initially unstable (i.e., neurons are drifting randomly relative to each other) but stabilises as a function of experience with the environment.

### Internal geometries differ across environments

CA1 neurons alter their spatial tuning when exposed to different environments (‘remapping’). Low spatial correlation and cross-decoding accuracy confirmed remapping between the two distinct environments (Supplemental Fig. 3A,B). Given that drift in spatial tuning over subsequent days in the same environment is characterised by a preserved internal geometry, we asked whether remapping upon novel context exposure similarly represents a geometry-preserving change. *G* angles were significantly more different between the familiar and novel environment than within the familiar track (Supplemental Fig. 3C). This analysis was conducted for day 20 of novel context exposure so as to capture population activity after stabilisation of the internal structure in the novel context. Distinct environments are thus represented by population activity with unique internal geometry.

### Coordinated drift allows spatial map recovery

The observations of stable population geometry and unstable spatial representations of individual neurons raise the question of how these phenomena can coexist. We asked whether this apparent contradiction can be resolved through similarity transformations (translation, scaling and rotation) of the neural activity. If drift indeed manifests in this fashion in CA1, we reasoned that it should be possible to correct (or “undo”) the drift by applying the inverse of these transformations to the population activity, and to thereby reveal a stable spatial map. We developed an approach to align the neural activities across days by translating, scaling and rotating population activity related to the geometric structure. Previous work that attempted to align neural activity across data sets relied on projections of the data into lower-dimensional manifolds and subsequent alignment within the reduced spaces^29–34^. However, with those approaches it is difficult to relate these transformations to the activity of individual neurons. Instead, in our approach, we applied dimensionality reduction techniques (demixed principal component analysis; dPCA) to identify the subspaces capturing the core structure of the task-related population activity. We then sought a transformation function that would align the subspaces in the full dimensional space from one (source) to subsequent (target) days (**Fig. 3A,B**). Importantly, this operation only allows rotation, translation and uniform scaling so as to preserve the internal geometry. After correcting the original high-dimensional dataset of the target days with the obtained transformation operation, we could recover highly stable spatial tuning functions over time, as evidenced by large spatial correlation values (**Fig. 3C,D**). Moreover, decoders trained on population activity on the source day were highly accurate in predicting the animal’s spatial position from calcium signals of the aligned target days compared to shuffled control data (**Fig. 3E**).

**Fig 3.**
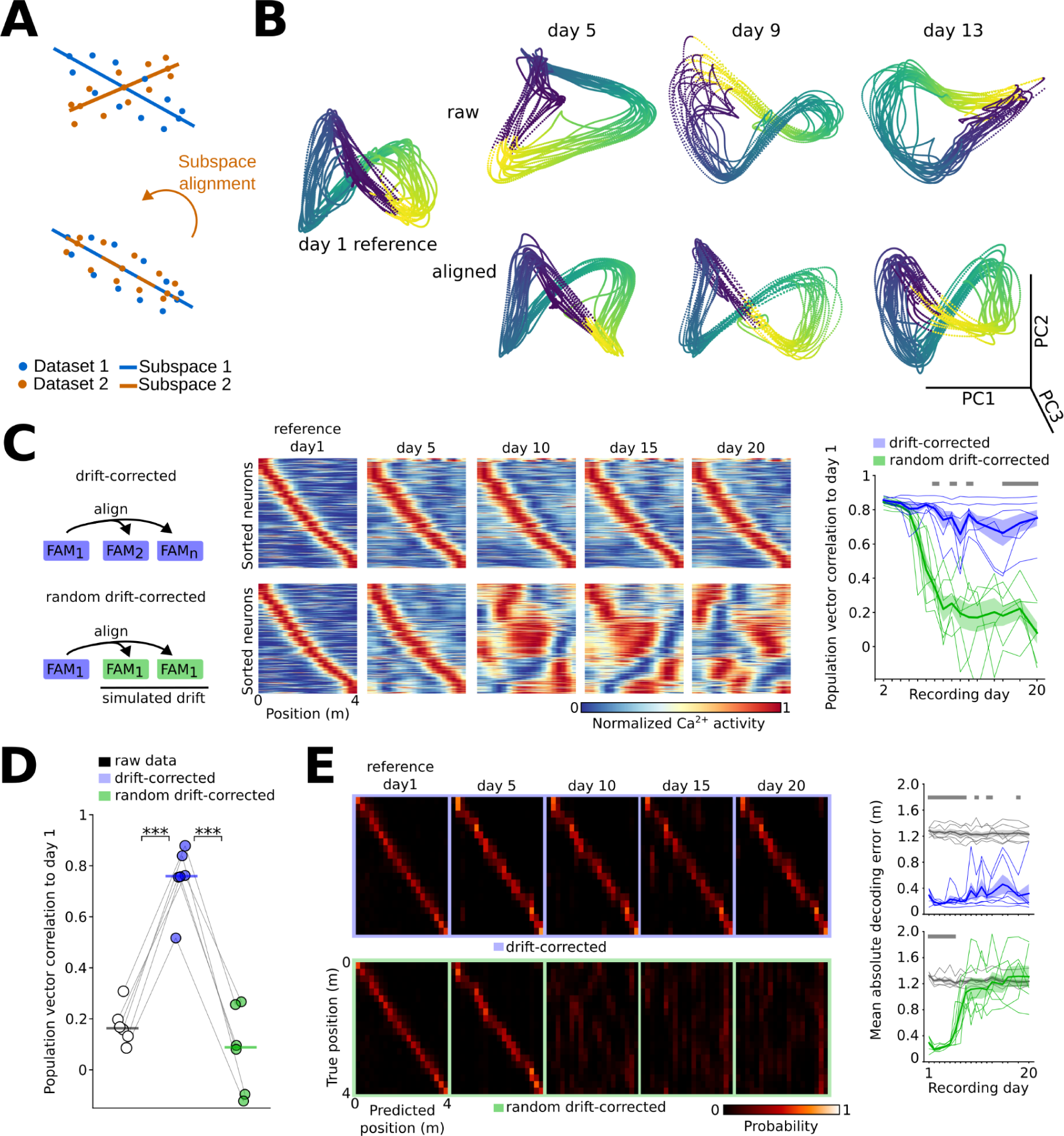
Representational drift can be corrected with geometric alignment. **A**. Illustration of geometric alignment. For each dataset, the neural data is translated to the origin and scaled to unit length. Then, the most significant subspaces are identified with dPCA. The datasets are aligned by rotating the subspace such that they end up overlapping. In this example, the data is low-dimensional (1 dimensional subspace is rotated in two dimensional space) with simple geometry. Aligning subspaces of high-dimensional geometric objects is a challenging task that only works if the objects share the same internal geometry. **B**. Data projected into the 3 first dPCA axes before and after subspace alignment, from an example mouse. **C**. Example of drift-corrected tuning functions for one mouse. Top: For day 1+n, the subspace in the familiar environment is rotated such that it best possible aligns with the subspace of day 1. Bottom: Same alignment procedure but for data with simulated uncoordinated (random) drift. Right: Population vector correlation for simulated data drops relative to aligned data with increasing time. t=6.39-29.07, p=0.024-10^-5^. **D.** Population vector correlation between the last imaging day and day 1 for raw data (white), drift-aligned data (blue) and aligned data with simulated drift (green). Raw vs. drift-corrected: t=-16.95, p=10^-5^; raw vs. random drift-corrected: t=1.96, p=0.318; drift-corrected vs. random drift-corrected: t=11.40, p=0.0003, 1-way repeated measures ANOVA followed by paired t-tests with Bonferroni correction. **E**. Decoding spatial position from all neurons after drift correction (top) and for simulated data with random drift (bottom). The confusion matrices show the probability of predicting each bin, averaged across all animals. Top right: Aligned data allows significant position decoding vs. shuffled data (grey). t=7.78-20.25, p=0.01-10^-5^. Bottom right: Predictions with simulated data quickly drop to chance level. t=6.39-29.07, p=0.024-10^-5^. C,E: Grey bars: significant differences, 2-way repeated measures ANOVAs followed by paired *t* or Wilcoxon tests with Bonferroni correction. Thick lines/shading: mean ± SEM, thin lines: individual mice (n=6).

To test whether effective alignment strictly requires a conserved internal geometry, we performed a series of control measurements. First, we applied the same alignment to data with simulated random drift (i.e., with non-conserved internal geometry). In this case, the recovered spatial correlation quickly dropped as a function of recording days, and decoding models could only correctly predict the animal’s location for a few future days, confirming that the temporal alignment can be performed with coordinated, but not with random drift (**Fig. 3C-E**). Second, we reasoned that in the presence of random remapping, alignment should not recover stable spatial tuning. Since CA1 pyramidal cells remap across environments in an uncoordinated manner, we do not expect their population activities to be alignable. Corrected tuning functions of the novel track indeed showed low spatial correlation to the familiar track, and decoding position on corrected novel track data with decoders trained on the familiar track produced large spatial errors (Supplemental Fig. 4A,B). Third, we used standard instead of demixed principal component analysis to extract a low-dimensional representation of population activity that captures the main covariance in the data (both task- and non-task related). Similar to our original approach, this procedure gave large spatial correlations and low decoding errors across days in the familiar environment, suggesting that alignment is robust towards the method of dimensionality reduction (Supplemental Fig. 4C). Finally, consistent spatial tuning could be extracted by aligning with neurons of varying levels of spatial information, indicating that the stable internal geometry does not strictly depend on neurons with strong place field characteristics (Supplemental Fig. 4D). Taken together, stable spatial tuning can be recovered by a similarity transformation of the original tuning functions.

### Coordinated drift is characterised by state space rotation and translation

Considering that distinct similarity transformations (rotation, translation and uniform scaling) are consistent with coordinated drift, we sought to understand the quantitative contribution of those transformations to the representational drift observed in CA1 (**Fig. 4A**). First, to quantify the degree of day-to-day rotation in state space, we calculated the subspace angles between the dPCA subspaces of day 1 and all subsequent days, and averaged angles across the principal axes. The results showed an increasing mean angular difference over days, suggesting a progressive rotation of the neuronal state space (**Fig. 4B**). This finding was consistent when we used other measures of rotation, including the minimum subspace angle, the trace of the alignment rotation matrix, and the Frobenius norm of the difference between the rotation matrix and the identity matrix (Supplemental Fig 5). Second, we assessed the extent of translation by taking the Frobenius norm of the difference between the centroid of the comparison day to day 1, scaled by the number of neurons in the dataset. This analysis revealed a significant gradual translation over time (**Fig. 4B**). Finally, we quantified scaling by the norm of the mean-centred data relative to day 1. In contrast to rotation or translation, scaling did not significantly change over days (**Fig. 4B**).

**Fig 4.**
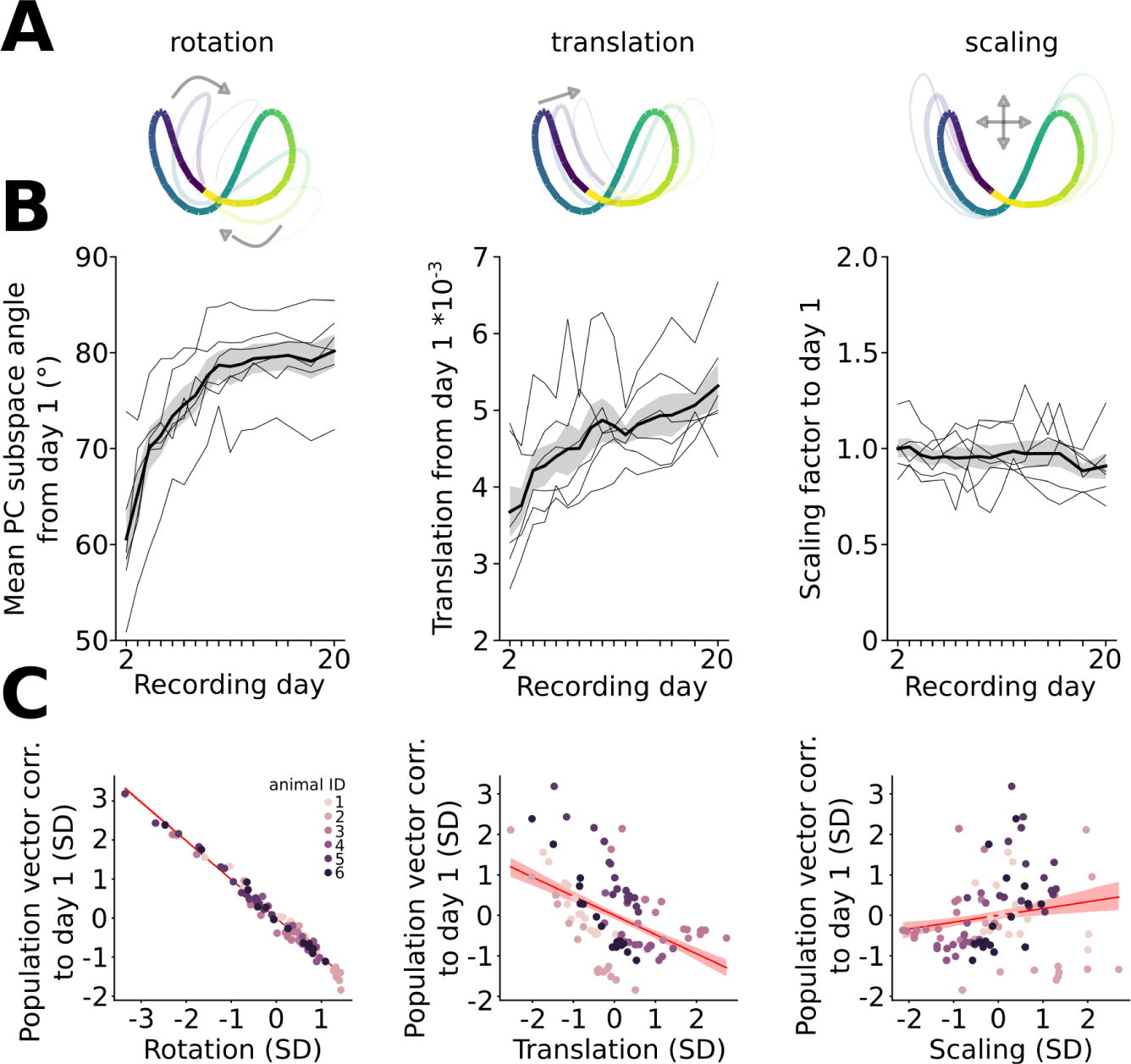
Representational drift is described by gradual rotation and translation in state space. **A.** Illustration of gradual similarity transformations with rotation, translation and uniform scaling. **B**. Rotation and translation relative to day 1 gradually increase over days (rotation: F=37.9, p=10-29, translation: F=15.8, p=10-4), with no change in uniform scaling (F=0.6, p=0.83, 1-way repeated measures ANOVAs). **C.** Linear mixed model estimating the relationship between drifting and each similarity transformation. For each transformation, the model controls for the effect of the other transformations. Rotation: β=-0.85, p=10-97. Translation: β=-0.18, p=10-4. Scaling: β=0.006, p=0.792. Thick lines/shading: mean ± SEM, thin lines: individual mice (n=6).

Given the gradual rotation and translation in state space across days, we investigated whether these similarity transformations could explain the representational drift observed on the spatial map. To isolate the effect of each transformation, we ran a linear mixed model with each transformation as predictors and drift as the outcome variable (quantified as the population vector correlation to day 1), sampled over all days and animals. The drift pattern could be well explained by the gradual rotation and translation in state space (**Fig. 4C**). In contrast, no relationship was found between drifting and scaling. Together, the results indicate that representational drift in CA1 can be explained by gradual rotation and translation in state space.

### Coordinated drift supports information recovery in downstream neurons

We have shown that the internal geometry of the population is stable over days and that stable spatial information can be recovered by aligning the geometric object through similarity transformations. This raises an important question: how can downstream neurons achieve such transformations in order to read a stable map?

The internal geometry contains a complete description of the encoded information, including what can be read by a downstream linear or nonlinear decoder^35–39^. This geometry defines how neural activity patterns for different spatial positions are organized in state space such that downstream networks can read out positions - for example, by using decision boundaries to separate these patterns^38^ (**Fig 5A**, Supplemental Fig 6A). To compensate for the upstream drift, a downstream reader could, in principle, apply a transformation to the incoming activity to bring the activity back to a stable orientation, as illustrated in Fig 3. However, this would require an upstream mechanism that modifies the incoming activity in real time, which seems biologically implausible. Alternatively, the downstream reader could compensate for the upstream drift by updating its weights, effectively rotating its decision boundary to align with the drifted activity (**Fig 5A**). Mathematically, these two mechanisms are equivalent, as applying the inverse of the upstream drift transformations, whether to the input or the downstream weights, cancels out the drift (Supplemental Fig 6A).

**Fig. 5:**
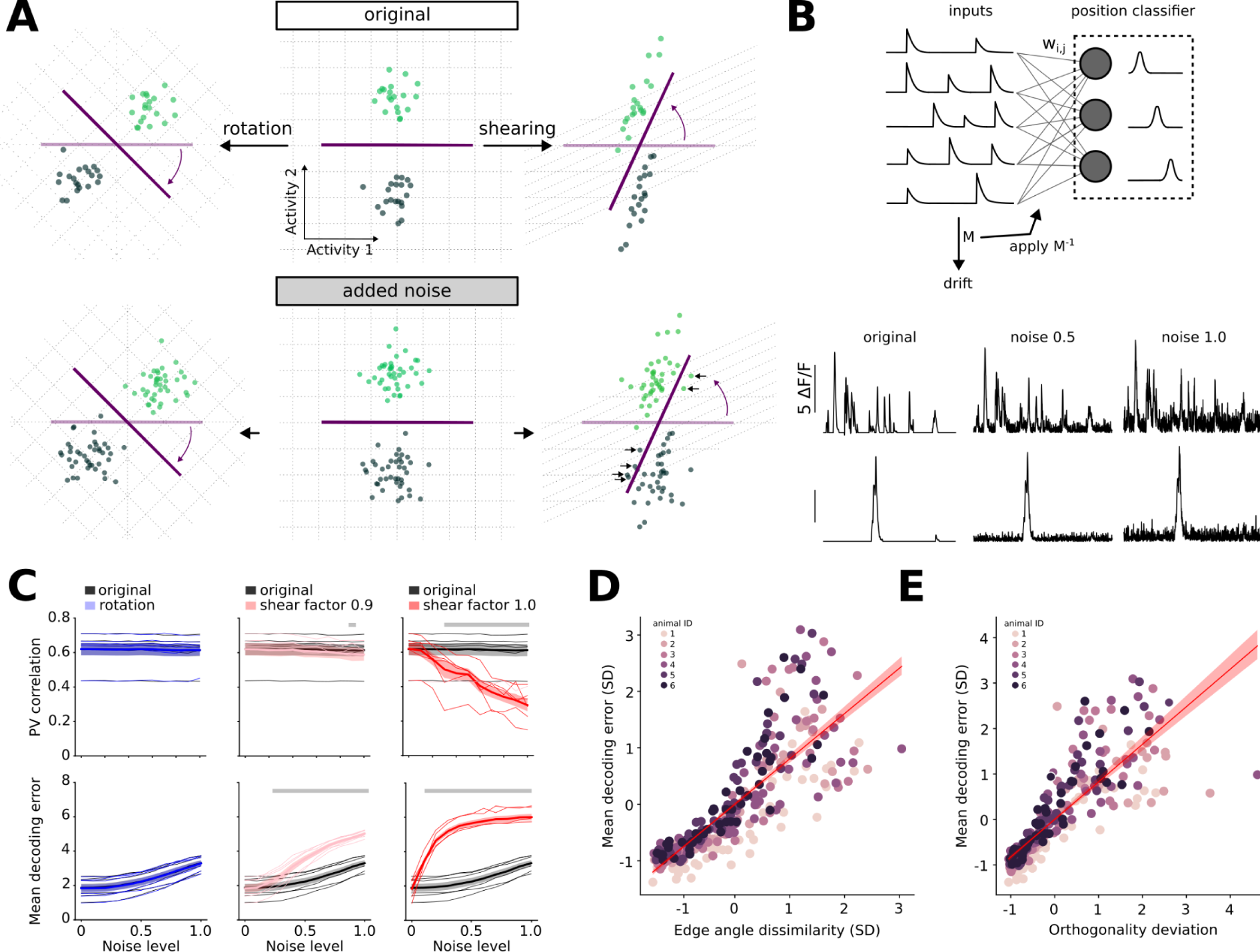
Coordinate drift supports information recovery in downstream neurons. **A.** Concept: A downstream linear decoder learns a decision boundary (purple) to classify positions (light and dark green) based on the activity of two neurons. A transformation applied to the input data moves the input relative to the decision boundary, which can be corrected by applying the inverse of the transformation to the decoding weights. Without noise, this works for both shape-preserving (rotation, left) and non-preserving transformations (shearing, right). Added noise results in classification errors in case of non-preserving transformations. **B.** Real inputs are fed into a positional decoder performing a linear classification with weights W_i,j_. The input is subjected to drift by applying a transformation matrix M, and the decoder is updated by applying the inverse of M to the weights. Bottom, artificial noise added at various levels to neuronal data. **C.** Top: Population vector (PV) correlation of the downstream neurons to the original state after drift correction. Noise decorrelates spatial tuning in downstream neurons for shear factors of 0.9 (t=11.64, p=0.0009) and 1.0 (t=5.42-10.01, p=0.032-0.0018) but not for rotation (F=0.45, p=0.531, effect of condition). Bottom: Mean absolute decoding error of the downstream network increases with noise in case of shearing (shear factor 0.9: t=7.79-19.22, p=0.006-10^-5^, shear factor 1.0: t=9.77-11.30, p=0.002-0-0009) but not rotation (F=0.01, p=0.942, effect of condition). Grey bars: significant differences, 2-way repeated measures ANOVAs followed by paired t-tests with Bonferroni correction **D.** Networks with larger edge angle dissimilarity show larger decoding errors. β=0.858, p=10^-176^. **E.** Networks with lower shape preservation (quantified as increasing orthogonality deviation) show larger decoding errors. β=0.817, p=10^-138^. D;E: linear mixed effect models. Thick lines/shading: mean ± SEM, thin lines: individual mice (n=6).

While it is theoretically possible to cancel out any upstream drift that has a defined inverse, we hypothesized that transformations that maintain the internal geometry of the population activity makes the system more robust to noise (such as input noise or inaccuracies in the weight updates; **Fig 5A**). To test this hypothesis, we simulated a downstream reader network that compensates for either geometrically preserved or distorted upstream drift under various noise levels (**Fig 5B**). Using the recorded CA1 activity of day 1 as input data, we modeled a population of downstream neurons with weights optimized such that each neuron is tuned to respond maximally to a specific location on the linear track. The upstream activity of day 1 was subjected to either a rotation (which preserves the geometry) or shear (which distorts the geometry), and the downstream neurons compensated for the drift by updating their weights using the inverse of the applied transformations. Both transformations allowed the downstream neurons to recover stable place tuning in the absence of added noise (**Fig 5C**). Furthermore, regardless of the drift, applying the inverse transformation restored the decoder’s ability to predict the actual position of the animal (**Fig 5C**). However, when we applied small Gaussian noise to the transformed input activity, the shear transformation performed significantly worse compared to the rotation, with increased difference as a function of noise level (**Fig 5C**).

To test whether this phenomenon generalizes beyond these specific transformations, we generated a diverse set of random well-conditioned transformation matrices with varying degrees of shape preservation (Methods) and examined their impact on downstream decoding. For each transformation applied to the upstream CA1 activity, the inverse transformation was applied to the downstream weights for drift compensation, followed by evaluating the decoding performance in the presence of noise. A clear relationship emerged from this analysis: the better a transformation maintained the internal geometry, the better the decoding performance under noise (**Fig 5D**, Supplemental Fig. 6B,C). Similarly, the performance was better when the transformations were closer to pure rotations (**Fig 5E**). These results could not be explained by the representational drift of the tuning responses, as drift per se was unrelated to the decoding performance after drift correction (Supplemental Fig. 6D). Overall, this suggests that shape-preserving transformations of CA1 activity might be advantageous for maintaining stable representation in downstream circuits despite biological noise.

## Discussion

We show that representational drift in mouse CA1 is not a random, uncoordinated process but can rather be described as a coordinated rotation and translation in neuronal state space. Drift in this form thus maintains the internal geometry of the population activity, allowing a downstream network to read stable information.

Our results expand upon previous accounts of conserved population activity across extended time periods. Earlier work in the neocortex^13,34^ and CA1^40^ has revealed long term stability by aligning low-dimensional projections of neural data across days. However, these studies did not address how this relates to representational drift of individual neurons. Furthermore, while previous studies have characterised the geometry in CA1 during spatial navigation^32,40–43^, a detailed quantification of the stability of the geometry across days was so far lacking. In the visual cortex of mice^13^, the correlations between population vectors are consistent over time, but it remains to be examined whether this corresponds to a stable internal geometry (or geometric shape) as quantified by a metric sensitive to shape but insensitive to rotation, translation and uniform scaling. To our knowledge, the stability of the internal geometry has to date only been investigated in such a way in the piriform^10^ and the prefrontal cortex^17^, but were found to be unstable in both regions across days. The CA1 circuitry stands in stark contrast to those brain areas as it is characterised by a stable geometry in the form of rotation and translation of the neuronal state space.

What mechanisms could give rise to the coordinated, geometry-preserving drift in CA1? One possibility is suggested by recent theoretical work showing that networks that combine feedforward inputs with lateral inhibition and Hebbian/anti-Hebbian plasticity optimize a geometry-preserving objective with multiple equivalent solutions^44^. Once in the optimal state, noisy synaptic updates drive the network to continuously explore geometrically equivalent solutions, resulting in translational and rotational coordinated drift. This mechanism aligns with CA1’s architecture of spatially tuned feedforward inputs from the entorhinal cortex and/or CA3^45,46^ showing Hebbian plasticity^47–49^, and local inhibition from parvalbumin-positive (PV) interneurons exhibiting anti-Hebbian plasticity onto pyramidal neurons^50^. Furthermore, the rapid translational and rotational population drift we observe aligns with CA1’s rapid synaptic turnover^51^. This model predicts that manipulating local PV interneurons should alter the population geometry, while modifying synaptic turnover should adjust the rate of translation and rotation.

A key question is whether the coordinated drift might serve a functional purpose. Inducing random shifts to the observed neuronal tuning functions resulted in an uncoordinated drift, demonstrating that the CA1 place map could, in principle, change in a random manner over days. Remapping between contexts is an example where this happens. It is possible that the circuit dynamics during repeated exposure to the same environment force the network to follow a coordinated change, which is not true across environments. Within the same environment, the geometry-preserving nature of drift might be advantageous for a downstream network through multiple mechanisms. From a population-coding perspective^52,53^, where computations are explained by population activity rather than tuning of individual neurons, trajectories through neuronal state space are a prominent candidate mechanism^54,55^. Here, coordinated drift preserves the shape of these trajectories despite changing single-cell activities. If the computation performed by the circuit resides in the trajectory’s geometry, coordinated drift will not alter the computational outcome of the circuit. Conversely, from a single-neuron perspective, we show that downstream circuits can theoretically compensate for upstream drift by adjusting their synaptic weights. The rigid transformations we observe in CA1 (rotation and translation) are particularly advantageous as they preserve distances between points in neural state space, making them robust to noise and straightforward to invert for the downstream decoder. While neurons are unlikely to have direct access to inverse rotation and translation matrices, biologically plausible learning mechanisms could achieve the same overall effect. Indeed, any mechanism that maintains stable downstream tuning despite transformation of upstream activity, must implement an inverse operation of that transformation. For instance, a downstream network updating weights to minimise the difference between their output and a stable internal^28^ or external^27^ reference signal, will effectively apply an inverse transformation. Further work is needed to identify the specific biological mechanism implementing this compensation in the brain.

The internal geometry associated with a familiar environment remains intact over time, at least over a time span of 20 days. A different environment, exemplified by the novel track in our study, shows a geometrically different structure. This opens the possibility that context information might be extracted from the population geometry. In other words, the internal geometry might serve as a barcode signal assigned to individual environments, facilitating discrimination between these arenas. This is equivalent to a pattern separation operation. Since population geometry within a given environment remains constant, individual neurons’ drift in their tuning functions might additionally facilitate the read-out of elapsed time from population activity. Consider a downstream network that compares the drift in the neurons’ tuning functions to a reference signal while tracking population geometry at the same time. This mechanism could keep track not only of which environment the agent is in but also for how long that particular environment has been visited.

## Methods

### Animals and surgical procedures

#### Animal experiments

All experiments involving animals were approved by the Regierungspräsidium Freiburg (license no. G20/137) in accordance with national legislation. For in vivo two-photon calcium imaging of CA1 pyramidal cells we used 6 Thy1-GCaMP6f mice (C57BL/6J-Tg(Thy1-GCaMP6f)GPS.5.17 DKim/J, 4 male, 2 female). We used mice of both sexes and aged 12-18 weeks at the beginning of the experiments. Mice were housed on a 12-h light–dark cycle in groups of 3 – 4 mice in a room maintained at a temperature of 21 °C (±1 °C) and relative humidity of 55% (±10%).

#### Surgery

All surgical procedures were performed in a stereotactic apparatus (Kopf instruments) under anesthesia with 1–2% isoflurane and analgesia using buprenorphine (0.1 mg/kg of body weight) and local application of lidocaine (2%). An eye lubricant ointment (Bepanthen, Bayer) was applied to protect corneal membranes during surgeries. The skin was first disinfected with 70% ethanol before the surgical incision, the dorsal surface of the skull was exposed and cleaned and a stainless-steel headplate (25×10×0.8 mm with 8 mm-wide central aperture) placed horizontally over the right hippocampus, centered on A/P = −2.0mm and M/L = - 1.9mm from Bregma and secured with dental cement (Super-Bond Universal Polymer Radiopaque, Catalyst V and Quick Monomer, Sun Medical). After curation of the cement a chronic cranial window was implanted. A craniotomy (diameter 3mm) was drilled at A/P − 2 mm, M/L - 1.9mm. Under continuous irrigation with chilled artificial cerebrospinal fluid, part of the somatosensory cortex and posterior parietal association cortex located above the hippocampus were progressively aspirated until the external capsule was exposed. The outer part of the external capsule was then gently aspired, leaving the inner capsule and the hippocampus intact. The imaging window implant consisted of a 3 mm-wide coverslip (CS-3R, Warner Instruments) glued to the bottom of a stainless-steel cannula (3 mm diameter, 1.4 mm height). This window was gently lowered into the craniotomy using forceps until the coverslip was sitting on the inner capsule. The implant was then fixed to the surrounding skull using cyanoacrylate. Post-operative analgesic treatment consisted of buprenorphine (2 injections of 0.1 mg/kg of body weight during daytime, overnight 1 mg/kg in drinking water) and carprofen (5 mg/kg of body weight, 12 h cycle) provided during 2 days after surgery. Mice were allowed to recover from surgery for at least 5 days before any further experiment.

### In vivo two-photon calcium imaging and behaviour

As previously described^3^, our custom virtual environment setup consisted of an air-supported polystyrene ball (20-cm diameter) attached on one side with a small metal axle (restraining the ball motion to the forward–backward direction). Ball movement was monitored using an optical sensor (G-500, Logitech) and translated into forward motion inside the virtual environment. The forward gain was set such that 4 m of distance traveled along the circumference of the ball equaled one full traversal of the linear track. When the mouse reached the end of the track, screens were blanked for 4–10 s before the mouse was “teleported” back to the start of the linear track. The virtual environment was displayed on four TFT monitors (19″ screen diagonal, Dell) arranged in a hexagonal arc around the mouse and placed ∼25 cm away from the head of the animal, thereby covering ∼260° of the horizontal and ∼60° of the vertical visual field of the mouse. The virtual environments were created and simulated using the open-source 3D rendering software Blender 2.79 (available at www.blender.org). The two different environments used in the present work consisted of distinct arrangements of textured walls, floors and 3D-rendered objects placed along the tracks sides. When the mouse reached any of the two rewarded sites (of which the positions were different for both environments; see Fig. 1A, 2A), 2 µl of oat milk were dispensed through a spout in front of the mouse. During training mice were first habituated to head fixation (for about 5-10 min per day) and subsequently allowed to explore a first (“familiar”) virtual environment for 10–30 min daily, with gradually increasing time spans over days. In parallel we set up food scheduling with a goal of ∼90% of the ad libitum body weight. Training in the familiar environment was maintained for 30–60 min daily until consistent reward licking and voluntary running were observed (i.e., at least 11 days of exposure to the familiar environment before any imaging session). After training we performed additional 3 baseline recordings to systematically habituate the mouse to all sounds of the imaging session.

#### Behavioral paradigm for imaging sessions

From the first day of imaging, mice were introduced to a novel environment, which had different visual cues and floor and wall textures, but had the same dimensions as the familiar environment. For each imaging session, mice alternatingly ran on the familiar and the novel tracks for a total of 40 runs per day. Runs were grouped by blocks of 5 (starting with the familiar one) for a total of 4 blocks and 20 recordings for each track. This recording procedure was repeated over 13 consecutive days and continued till day 20 but with breaks on days 14, 17 and 19. During each recording the exact same field of view (and therefore the same population of CA1 pyramidal cells) was imaged. We performed this experiment in 6 Thy1-GCaMP6f mice in which we imaged a total of 5043 CA1 pyramidal cells (mean number of cells imaged per mouse ± SEM: 840.5 ± 93.8).

#### Reward-related licking behavior

Licking was monitored using an infrared optical lick detector placed in front of the metal lick spout dispensing the reward. The reward zones were defined as the zone spanning 10 cm before and 40 cm after the reward sites, excluding the first and last 10 cm of the track. The remaining track was considered as outside of the reward zones. Thus, the ratio of licking between the inside and outside of the reward zones was computed as the mean lick rate in the reward zones divided by the mean lick rate on the remaining track. Likewise the ratio of the running velocity of the mouse inside and outside of the reward site was calculated. Only running periods(≥ 5 cm/s) were considered.

#### In vivo two-photon calcium imaging

In vivo calcium imaging was performed using a resonant/galvo high-speed laser scanning 2-photon microscope (Neurolabware) through a 16x objective (Nikon, 0.8 N.A., 3 mmWD) with a frame rate of 15.5 Hz and using a single plane for imaging. GCaMP6s was excited at 930 nm with a femtosecond-pulsed two-photon laser (InSight DS+, Spectra-Physics). To block ambient light from reaching the photodetectors, the animal’s head plate was attached to the bottom of an opaque imaging chamber before each experiment, and the mouse was then affixed to the virtual environment setup using this head chamber. A ring of black foam was placed between the imaging chamber and the microscope objective, and a metallic collar surrounding the objective was sitting on the imaging chamber, blocking any remaining stray light. Laser power and photomultiplier (PMT) detectors (Hamamatsu H11706-40 GaAsP) were compensated appropriately for each imaging session, ensuring consistent recording conditions. Data was acquired using the Scanbox software (Neurolabware).

### Extraction of calcium signals and data processing

The processing of all raw calcium data (pooled for all days) was done using the Python-based toolbox Suite2p^56^, a free automated pipeline for processing two-photon calcium imaging recordings (available at www.github.com/Mouseland/suite2p). Briefly, Suite2p first aligns all frames of a calcium movie using two-dimensional rigid registration based on regularized phase correlation, subpixel interpolation, and kriging. This toolbox then allows visual inspection of the registered movie. In all 6 included datasets a consistent alignment was achieved over the entire course of the experiment (i.e., 17 recording days spanning 20 days). Suite2p then performs automated cell detection by computing a low-dimensional decomposition of the data, which is used to run a clustering algorithm that finds regions of interest (ROIs) based on the correlation of the pixels inside them. All ROIs were manually curated to ensure the most accurate selection of pyramidal cells. ROIs that showed activity patterns of multiple cells or composed cell compartments (axons, dendrites) were discarded. Furthermore we verified that segmented entities were clearly visible throughout the entire experiment.

From the extracted raw fluorescence per ROI significant calcium transients were identified as previously described^3,56,57^. Imaging was performed ∼100–200 µm under the window, in the middle of the proximo-distal CA1 axis with each FOV containing deep and superficial pyramidal cells, thus our data included cells imaged along the entire radial axis of the pyramidal cell layer. We restricted our analyses to periods with a running speed of at least 5 cm*s^−1^. In brief, calcium traces were corrected for slow changes in fluorescence by subtracting the 8th percentile value of the fluorescence-value distribution in a window of 20 s around each time point from the raw fluorescence trace. We obtained an initial estimate on baseline fluorescence by calculating the mean and standard deviation (s.d.) of all points of the fluorescence signal that did not exceed 2.3 s.d. of the total signal. We then divided the raw fluorescence trace by this value to obtain the ΔF/F trace. This trace was used to determine the parameters for transient detection that yielded a false positive rate (defined as the ratio of negative to positive oriented transients) <5% and extracted all significant transients from the raw ΔF/F trace. Definitive values for baseline fluorescence and baseline s.d. were calculated from all points of the trace that did not contain significant transients. A transient mask was created, and for further analysis, all values of this ΔF/F trace that did not contain significant calcium transients were set to zero in order to improve the signal-to-noise ratio. Using this method, ΔF/F is expressed in units of s.d. (the standard deviation of the baseline fluorescence).

### Analysis of spatial tuning functions across days

For each session, the z-scored ΔF/F trace of each neuron was averaged within spatial bins based on the animal’s position along the linear track. The linear track was divided into 20 bins, and mean activity of the neurons were calculated for each bin. The tuning functions were sorted either by a reference session (such as day 1) or within each session. For visualization of the spatial maps, the binned activity was normalised per neuron so that the bin with the largest values was scaled to 1. Furthermore, inactive neurons (<2% of active frames) within a session were not included when visualizing the spatial map. However, all neurons were included for sorting the maps and downstream analysis steps.

### Spatial information

As previously described^58^, the spatial information (*SI*) for each neuron was calculated as

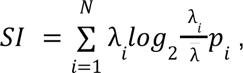

in which λ_*i*_ is the mean activity of the neuron in the ith bin, λ is the average calcium activity over the entire track, *p*_*i*_ is the occupation probability of the ith bin, and *N* is the number of bins on the track (*N*=20).

### Decoding spatial position

For decoding spatial position (20 digitised positions) of animals from neuronal activity, we used a Gaussian Naive Bayes classifier implemented with sklearn.naive_bayes.GaussianNB. Prior to training, the calcium signals were z-scored to ensure the features were on a comparable scale. The model was trained on the first half session and tested on the remaining half session of day 1. This provided a baseline measure of the model’s performance. The model was then used to predict spatial position on data from subsequent sessions (days 2 to 20) to evaluate how well the spatial representations generalised to new days. The mean absolute error (MAE) was used to evaluate the model’s performance. To assess the significance of the decoding performance, we statistically compared the data against a shuffle control. Before generating train and test data, we independently circularly shifted the calcium signal of each neuron by a random time step. This way of shuffling preserves the overall activity patterns but disrupts the temporal relationship between individual neurons.

### Quantifying representational drift

Representational drift of the spatial tuning functions was calculated by comparing the tuning functions (see *Analysis of spatial tuning functions across days*) between the first day and all subsequent days. As a measure of overall drift across all neurons between two sessions, we flattened the spatial maps and calculated the Pearson’s correlation coefficient. Within-day stability was estimated using a similar approach but correlating odd and even trials. We also performed a cross-decoding approach to test generalization of spatial representations across days (See *Decoding spatial position*). To estimate how much an individual neuron drifted, we computed the spatial map correlation for each neuron individually. These correlations were averaged across days for each neuron separately to measure overall stability of individual neurons. Since some neurons were inactive in some sessions, we did not consider inactive days for this analysis. Furthermore, stability was not computed for neurons with less than 4 active sessions.

### Dimensionality reduction

To extract low-dimensional subspaces from the calcium activity, we used two complementary approaches. Standard principal component analysis (PCA) was applied to get insights into the overall activity structure, while demixed principal component analysis (dPCA^59^) was used to focus on only task specific neural dynamics. For each session *s*, the z-scored calcium activity was represented as *X*_*s*_ ∈ ℝ^*T*×*N*^, where *T* is the number of timepoints and *N* is the number of neurons. Both methods work by finding projection matrices that transform the high-dimensional neural activity into a lower-dimensional space. The key difference is that while PCA captures directions of maximal variance in the data, dPCA isolates neural activity that is specifically related to the spatial positions of the animal. The projection is computed as:

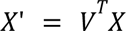

Where *X* represents the calcium activity and *V* ∈ ℝ^*N*×*k*^ is the projection matrix. For both methods, we projected the data into the first *k* = 6 components.

### Simulating drift

To compare our experimental data suggesting coordinated CA1 drift to a hypothetical example of uncoordinated drift, we implemented simulations that induced uncoordinated drift between neurons while keeping the overall spatial tuning and firing rate constant. For each neuron independently, we included two forms of drift: spatial tuning remapping and firing rate drift. Spatial tuning remapping was modeled as a probabilistic remapping process. On each day, each neuron had 30% probability of shifting their place field to a random new location. The shift was applied in a circular manner in order to maintain the shape of the place fields while changing the location of the field. The same shifts were applied proportionally across all individual runs to maintain consistent spatial tuning.

The firing rate drift was modelled as a multiplicative random walk model. On each day, each neuron had a 30% probability to undergo amplitude change. For neurons selected to change amplitude, amplitude magnitude change factors were drawn from a log-normal distribution to ensure a skewed distribution with mostly small changes and occasionally larger changes. This was implemented by generating random numbers ∈ from *N*(0, σ^2^) with σ = 0.5, followed by an exponential transform corrected to have an expected value of 1:

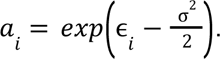

The magnitude change factor *a*_*i*_ was applied multiplicatively to the activity trace for neuron *i* for each run on the linear track. This process ensured that neurons had an equal chance of increasing or decreasing their amplitude while keeping the mean amplitude across the population stable over time. We also tested other drift parameters with different drift probabilities (10%, 30%, 60%), amplitude change probabilities (10%, 30%, 60%) and amplitude magnitude change factors (σ = 0.25, 0.5, 0.75).

### Analysis of internal population geometry

To investigate the stability of the internal geometry of the population responses across days, we applied three complementary approaches: edge angle analysis, subspace angles analysis, and persistent homology. These methods allowed us to quantify changes in population geometry while being insensitive to shape-preserving translation, rotation or uniform scaling in the neural space.

#### Edge angles

In the edge angle analysis^10^, we connected edges between every 3 positions in the spatially binned map for each day. Edges were connected by subtracting a third positional vector from the two other vectors. The angle between a pair of connected edges were computed as follows:

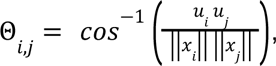

where Θ_*i*,*j*_ is the angle between vector *i* and *j*. Edge angle analysis has previously been successful in analysing changes in internal population activity across days in piriform^10^ and prefrontal cortex^17^.

#### Subspace angles

As a complementary approach, we quantified changes in local subspace angles across days. We defined subspaces by grouping population vectors from consecutive positional bins. Thus, for each pair of spatial bin *i* and *j*, we constructed spaces *S*_*i*_ and *S*_*j*_ by taking the bins at position *i* to *i* + 1 and *j* to *j* + 1. To compute the principal angles between these subspaces, we first computed orthonormal bases *Q*_*S_i_*_ and *Q*_*S_j_*_ for the subspaces *S*_*i*_ and *S*_*j*_ using QR decomposition (from *scipy* module *linalg.orth*). We then computed the matrix:

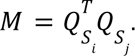

The cosines of the principal angles between these subspaces were extracted from the singular values in:

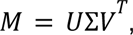

where Σ is the diagonal elements containing the singular values σ_*k*_. Then, the principal angles are calculated as:

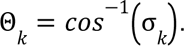

#### Persistent homology

To capture higher-order topological features of the population activity, we used persistent homology. This method identifies topological features such as connected points, holes and higher dimensional voids that persist across multiple scales. The method works by creating a sequence of simplicial complexes by progressively connecting nearby points based on a distance threshold that increases from 0 to a maximum value. As the threshold increases, some features emerge (birth time) or disappear (death time). The persistence of the features are described as the difference between the death time and the birth time. Using the *Ripser* algorithm^60^, we computed persistence diagrams for dimensions up to 4. These diagrams represented the birth and death times of the topological features in the data. Persistent cohomology had previously been used to describe the topology of head direction^40,61,62^-, place-^63,64^ and grid cells^65–67^.

#### Dissimilarity metrics

To assess the long-term stability of the internal geometry, we computed dissimilarity metrics for each method. For edge angle and subspace angle analysis, we computed the Frobenius norm on the difference between the angle matrices from days. For a pair of angle matrices *A^p^* and *A^q^*, corresponding to day *p* and *q*, respectively, the Frobenius norm was calculated as

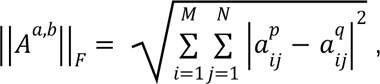

where *N* is the number of starting positions (spatial bins for edge angles and subspaces for subspace angles), and *M* is the number of possible angles to measure 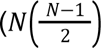 for edge angles and 2(*N* − 1) for subspace angles). A Frobenius norm of 0 indicates identical angle matrices, while higher values indicate increasing dissimilarities.

For persistent homology, the stability was assessed by comparing persistent diagrams from day *p* and *q* by calculating the Wasserstein distance averaged across all dimensions:

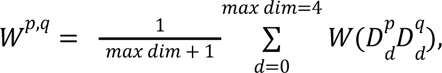

where *W* denotes the Wasserstein distance between persistent diagrams *D^*p*^*_*d*_ and *D*^*q*^_*d*_ in dimension *d* calculated with *persim.wasserstein*.

#### Matrix normalization

To enable comparison of within-day and across day variability, we normalised the matrix dissimilarity matrices as previously described^10^. For edge angle and subspace angle analysis, the matrix dissimilarity matrix ||*A^a,b^*||_*F*_ was normalized in the following way:

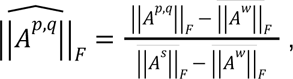

where 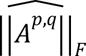 is the normalised dissimilarity measure, 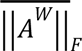 is the within-day Frobenius norm computed between odd and even runs and averaged across all days, and 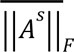 is the Frobenius norm between shuffled data on the first and last day. In this normalised metric, 0 corresponds to average within-day dissimilarity while 1 unit is equivalent to the dissimilarity of the shuffled data. For the persistent diagrams, since the Wasserstein distance were smaller between shuffled data than any real data, we simplified the normalization for persistent homology dissimilarities to:

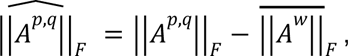

such that 0 corresponds to the average within-day dissimilarity, without any upper bound.

#### Analysis variations

We performed this analysis in different variations. Forward direction analysis compared day 1 with all days up to 20, while reverse direction analysis compared day 20 with all days back to day 1. This analysis was performed separately on data from familiar and novel environments, and on simulated drift data (see *Simulating drift*). For some additional analysis, we compared the geometry across the two environments (familiar and novel), relative to within the same environment. In this case, the matrix dissimilarities were calculated between familiar and novel on day 20 for across environments, and between familiar day 17 and 20 for within-environment. Within- and between environment matrix dissimilarities were normalised with same data, such that 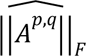 is the within-day Frobenius norm averaged over the two context, and 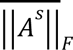 is the Frobenius norm between shuffled data on day 17 of familiar environment and day 20 of novel environment. We also reanalysed the data by focusing on only task-related subspaces, using PCA and dPCA (see section *Dimensionality reduction*).

### Recovering spatial tuning with geometric alignment

If the internal geometry of the neural population activity in CA1 is stable over days, we hypothesised that it should be possible to recover a stable spatial map by aligning the underlying geometric structure across days. To recover the spatial maps, we developed a method to align the full-dimensional neural activities across days by translating, scaling and rotating the data according to subspaces related to the geometric structure.

To align the neural activity across days, we first translated each dataset into a common reference frame by centering the data into their origins. We then globally scaled the centered data into unit length by dividing all elements by the Frobenius norm. We then applied PCA or dPCA to the neural data *X* ∈ ℝ*^T^*^×*X*^, where *T* represents the number of timepoints and *N* is the number of neurons. From this decomposition, we extracted the first *k* = 6 principal components, forming the matrices *P*_1_ ∈ ℝ^*N*×*k*^ and *P*_2_ ∈ ℝ^*N*×*k*^, which correspond to the principal axes of the reference day and the day to be aligned, respectively. These axes define low-dimensional subspaces capturing the core geometric structure of the population activity.

The next step of the alignment process involves rotating the subspace spanned by the principal axes of the first day so that it best overlaps with the subspace spanned by the reference day. Using a weighted orthogonal procrustes transformation, we searched for a rotation matrix *R* ∈ ℝ^*k*×*k*^ that minimizes the weighted Frobenius norm of the difference between the aligned subspaces:

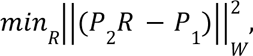

where ||·||_*W*_ denotes the weighted Frobenius norm, and *W* is a diagonal matrix of *k* weight:

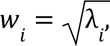

with λ_*i*_ being the explained variance ratios from PCA or dPCA. These weights were incorporated by scaling the principal axes:

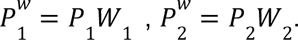

The optimal rotation matrix *R* is obtained using singular values decomposition (SVD), which decomposes the weighted cross-covariance matrix as:

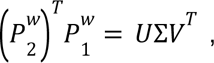

where *U* and *V* are orthogonal matrices, and Σ is a diagonal matrix containing the singular values. The optimal rotation matrix *R* is given by:

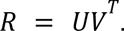

To apply the rotation to the original high-dimensional space, we expanded *R* to form a transformation matrix *R_high_* ∈ ℝ^*N*×*N*^ as follows:

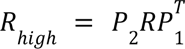

The high-dimensional transformation matrix *R_high_* was then applied to the neural activity *X*_2_ from the day to be aligned:

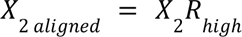

In effect, the whole transformation rotates the task-related subspace of *X*_2_ such that it best possible overlaps with the task-related subspace of *X*_1_, without distorting the internal population geometry.

### Quantification of similarity transformations

To assess how the population geometry translates, scales and rotates across days, we quantified these similarity transformations for each day relative to the first day. This way, we could determine whether the location, size and orientation of the internal population geometry became progressively more different from day 1. These measures were captured as part of the alignment of the neural data across days (see *Recovering spatial tuning with geometric alignment*). For each day’s dataset, we calculated the centroid of its geometric object by averaging the neural activity across all timepoints. This value corresponds to the overall translation of the high dimensional data relative to the origin. To quantify the translation between two days, we computed the Euclidean distance between their centroids, normalized by the number of neurons *N*:

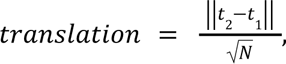

where *t*_1_and *t*_2_are the centroids of day 1 and day 2, respectively. The normalization accounts for differences in the number of neurons between animals. To quantify scaling, we computed the magnitude of the centered neural activity as:

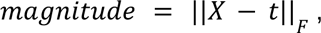

where *X* is the neural activity and *F* denotes the Frobenius norm. To get the change in scale across two sessions, we calculated a scaling factor as a ratio of their magnitudes:

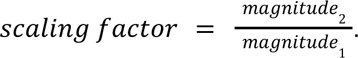

After centering and scaling the data, we aligned the neural data across days using the weighted orthogonal procrustes transformation on the principal axes as described in *Recovering spatial tuning with geometric alignment*. To quantify how much the data rotated between two days, we quantified the principal angles between the subspaces spanned by their principal axes. The *n* angles from the *n* principal axes were converted from radian to degrees and averaged to give a description of overall rotation. We also used three additional measures to quantify the degree of rotation. First, we calculated the minimum instead of the mean subspace angle among the *n* principal axes. Secondly, we computed the trace of the rotation matrix used to align the neural data scaled by the number of neurons:

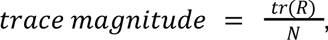

where R is the rotation matrix and N is the number of neurons. Finally, we quantified the deviation of the rotation from the identity matrix using the Frobenius norm:

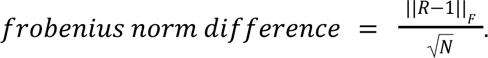

### Simulating upstream drift and downstream weight corrections

We modeled a downstream population of 20 neurons receiving inputs from the recorded CA1 neurons. Each downstream neuron was fitted to be maximally tuned to one of 20 digitized positions along the track. The downstream weights *W* ∈ ℝ^*N*×*P*^ were optimized using ridge regression:

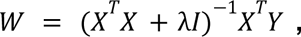

where *X* ∈ ℝ^*T*×*N*^ represents the calcium activity with *T* timepoints and *N* neurons, *Y* ∈ ℝ^*T*×*P*^ is a one-hot encoded position matrix with *P* spatial positions, and λ = 10 is a regularization parameter. The predicted position was estimated by taking the maximum response across the downstream population. We used half of the data for training, and the remaining half for testing.

To simulate upstream drift, we applied one of three types of transformations to the recorded activity. First, to simulate shape-preserving rigid transformation we applied random orthogonal matrices *M* ∈ ℝ^*N*×*N*^ from the special orthogonal group SO(n) (using *scipy.stats.special_ortho_group)*. Second, shape-distorting transformations were simulated by applying shears constructed as

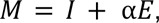

where *I* is the identity matrix, *E* has ones on the first upper diagonal and α is a scale factor of 0.9 or 1. Finally, for more general transformations of varying orthogonality (or shape-preservation), we generated random transformation matrices as

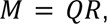

where *Q* is a random orthonormal matrix from SO(n) and R is a symmetric positive-definite matrix with controlled singular value distribution:

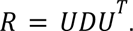

Here, *U* is another random orthonormal matrix, and *D* is a diagonal matrix. To perturb the orthogonality of the matrix, the diagonal entries of *D* were set to 1 in most dimensions but modified in 30% of randomly selected dimensions by multiplying with *exp*(ε), where ε ∼ *N*(0, σ^2^) with σ randomly sampled between 0 and 5. Finally, the determinant of *R* was scaled to 1, and the matrix was retained only if the condition number was below 10. To ensure a variety of different perturbations, we constructed 50 matrices *M* per animal.

For each transformation matrix *M*, we simulated the upstream drift by applying *M* to the test data as

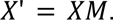

To let the downstream network compensate for the drift, we updated the downstream weights using the inverse transformation:

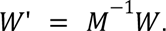

To test how robust the transformations were to noise, we added independent Gaussian noise to the drift transformed data:

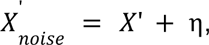

where η ∼ *N*(0, σ^2^). We tried different noise levels σ ranging from 0 (no noise) to 1. We then fed these noise added inputs *X_noise_* to the drift corrected downstream network with weights *W*’ to obtain downstream tuning functions and position predictions.

The performance against noise was evaluated in two ways. First, we computed the mean absolute error (MAE) between the true and predicted positions from the downstream decoder. Second, we quantified the stability of spatial tuning in individual downstream neurons by calculating the population vector correlation (see *Quantifying representational drift*) between the spatial maps before and after applying the transformations and noise.

The orthogonality of the transformation matrices were quantified as:

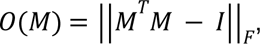

where ||·||_*F*_ is the Frobenius norm. A value of *O*(*M*) = 0 indicates a perfect rotation, whereas larger values correspond to increased geometric distortions. In addition, we quantified how well each transformation preserved the internal geometry of the upstream activity using edge angle, subspace angle and persistent homology analysis on the transformed neural data (see *analysis of internal population geometry*).

### Data analysis and statistics

All data analysis and statistical tests were conducted with custom written Python scripts using open-source libraries including numpy (version 2.33.3), scipy (1.10.1), pandas (1.4.2), matplotlib (3.7.2), scikit-learn (1.3.0), statsmodels (0.14.0), pingouin (0.5.3), h5py (1.12.1), dpca (1.0.5), ripser (0.6.8) and persim (0.3.7). For statistical comparisons involving multiple time points, we implemented repeated measures ANOVA (one-way for a single group and two-way for more groups). Statistically significant effects were followed up with post-hoc pairwise comparisons. Prior to pairwise comparisons, we assessed the normality of the data using the Shapiro-Wilk test. Paired t-tests were used for normally distributed data, while non-parametric Mann-Whitney U tests were conducted for non-normal distributions. In case of multiple comparisons, all resultings p-values were corrected with Bonferroni correction. Linear mixed effects models (mixedlm in statsmodels) were used for correlational analysis in which neural data was pooled across multiple animals. By treating the variables of interest as fixed effects and animal ID as random effect, this approach allowed us to account for variability between animals. For these analysis, we report standardized coefficients (β) and the corresponding p-values for the fixed effects. Throughout our analysis, the statistical tests were 2-sided. Data are expressed as mean ± SEM.

## Acknowledgements

This work was supported by the Else Kröner-Fresenius-Stiftung (grant A173_2019 to J.-F.S.), the German Research Foundation (FOR5159, grants to J.-F.S. (SA 3609/2-1) and to M.B. (BA 1582/16-1), BA 1582/15-1 to M.B., the CRC/TRR384 IN-CODE to M.B. and J.-F.S)), the Hans A. Krebs Medical Scientist Programme (to A.K.) and the ERC-AdG (787450 to M.B.).

## Author information

### Contributions

O.C.S., A.K., M.B., and J.-F.S. conceptualised the study. O.C.S. designed and performed analyses. A.K. performed experiments and calcium signal extraction. M.B. and J.-F.S. performed supervision and funding acquisition, O.C.S and J.-F.S. wrote the manuscript with contributions from all authors. All authors edited the manuscript.

## Supplemental figures and tables

**Supplemental Fig. 1:**
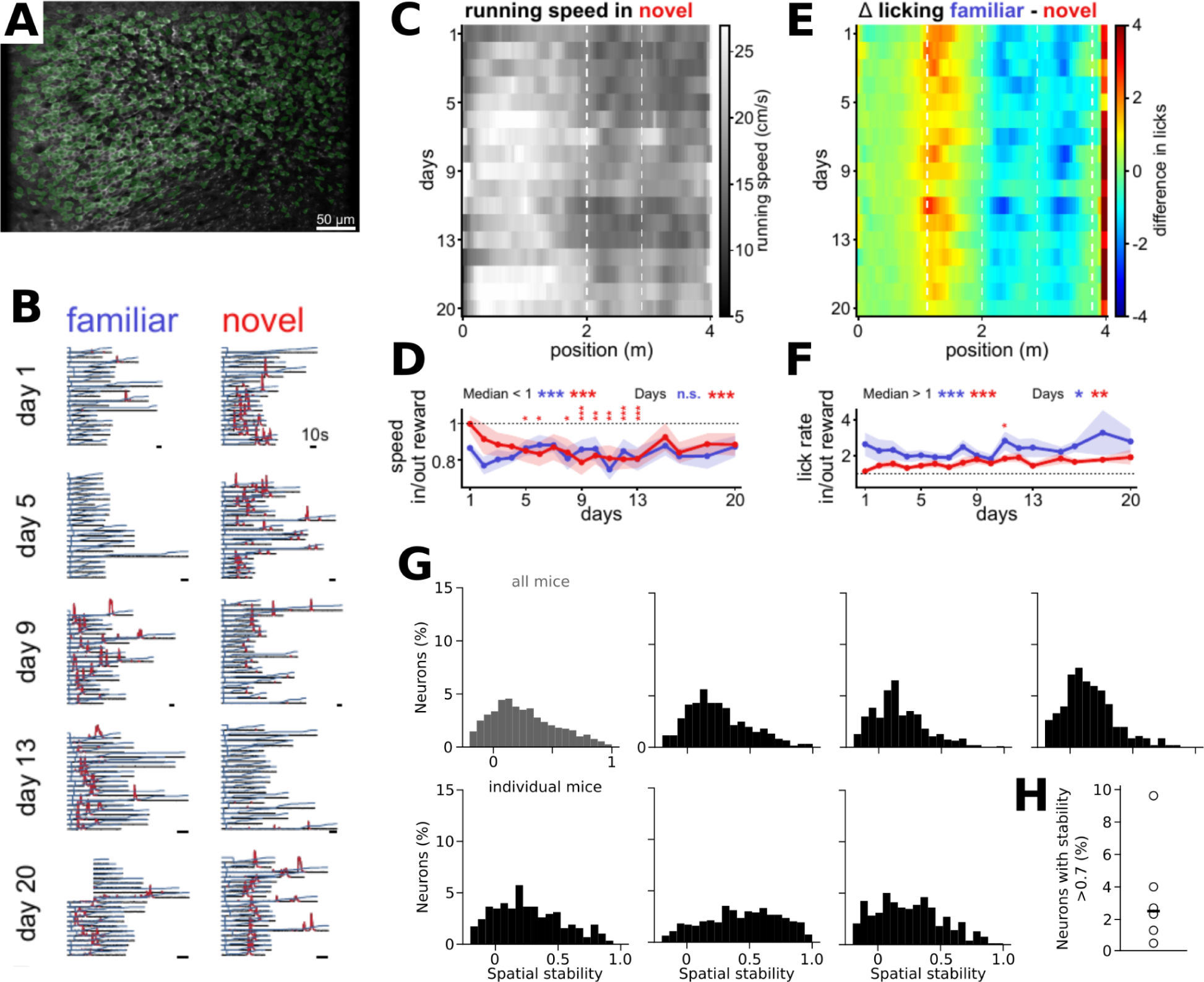
**A.** Mean image of the field of view in CA1 pyramidal cell layer after motion correction of all pooled 4 m runs acquired over 20 days (17 recording days). Identified ROIs of individual cells are overlaid in green. **B.** Exemplifying CA1 pyramidal cell activity over time in familiar and novel context on different days. In baseline-corrected fluorescence traces (black) acquired while the mouse was running along the 4 m track (position in blue) significant transients (red) are detected. **C.** Average running speed along the linear track in context novel (mice=6) over 20 days (only periods of movement ≥ 5cm/s considered). **D.** Ratio of running speed close and distant to the reward site is in both, novel (red) and familiar (blue) context below 1 (mice and time points pooled for one-sample signed rank test for hypothesized median = 1, median(fam) = 0.860, median(nov) = 0.877 fam/nov: p < 0.001), indicating that the mice slow down around reward sites. In familiar, this behavior is stable over time (1-way ANOVA repeated measures, p(days) = 0.233) while on the first day in the novel context they do not modulate their running speed around the reward site (1-way ANOVA repeated measures, main effect of days p < 0.001, stars indicate Tukey’s posthoc comparison to day1, all other day-by-day comparisons p > 0.05, mice = 6, mean ± sem shown). **E.** Heatmap showing the difference in licking rate between familiar and novel context averaged across mice for each imaging day. Mice show context specific increase of licking around reward sites (white dashed lines, blue/red drops). **F.** Ratio of lick rate close and distant to the reward site is in both contexts above 1 (mice and time points pooled, one-sample signed rank test for hypothesized median =1, median(fam) = 1.925, median(nov) = 1.477, fam/nov: p < 0.001), indicating that mice lick more within the reward zones. Despite some fluctuation over days (repeated measures on ranks, fam: p(days) = 0.021, nov: p(days) = 0.006, Tukey’s posthoc day-by-day all comparisons p > 0.05 but day 1vs11 in novel p=0.039) there is no systematic change in licking behavior with time (mice = 6, mean ± sem shown). **G.** Distribution of spatial stability (i.e., spatial correlation of the neurons’ tuning functions between day 1 and 20) pooled over all mice (grey) and for each mouse separately (black). **H.** Percentage of neurons with stable place field location (spatial correlation between day 20 and day 1 >0.7).

**Supplemental Fig. 2:**
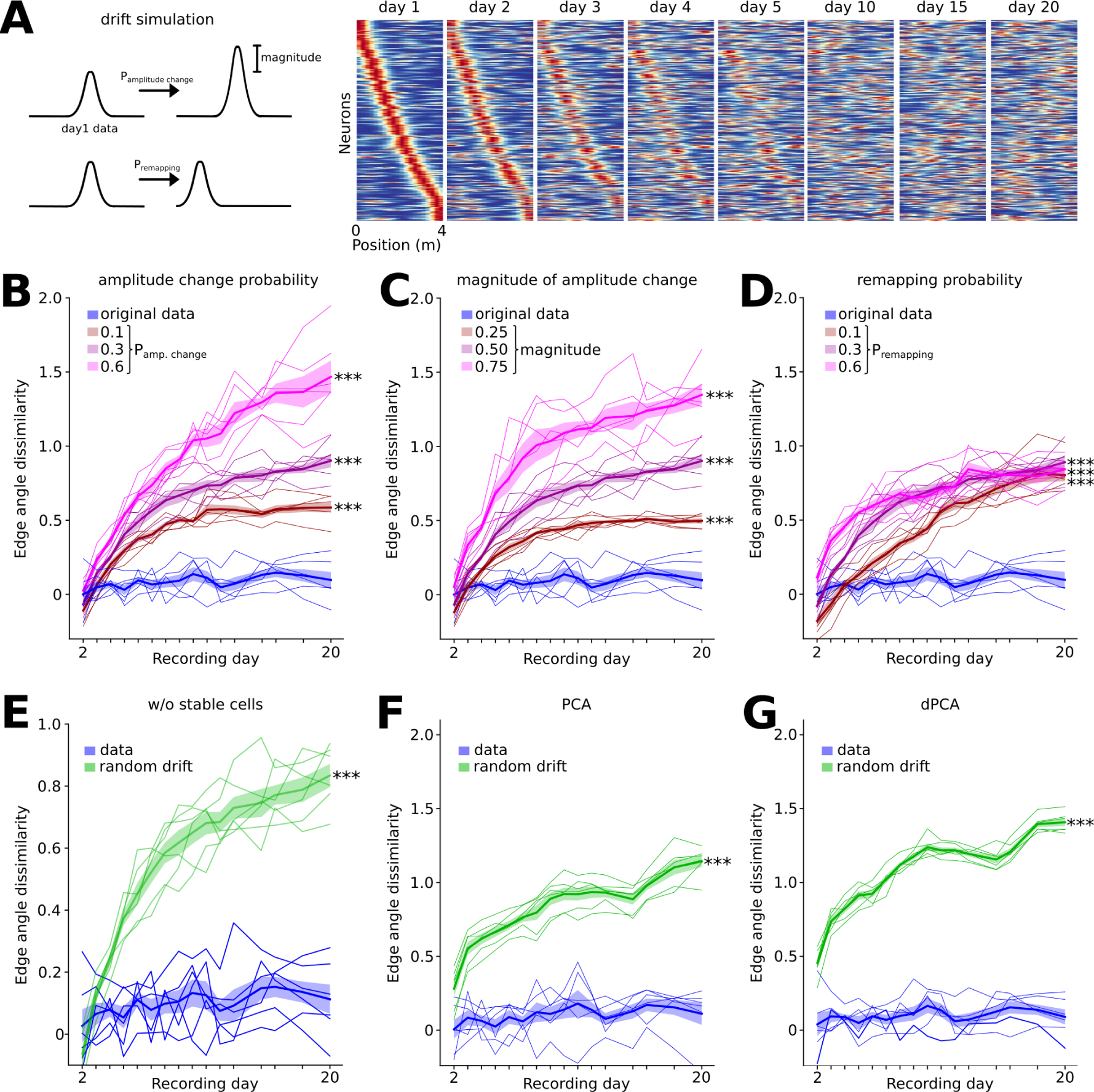
Geometry dissimilarity measures for different simulated drift levels and spatial stability. **A.** Left: Schematic of drift simulation. Amplitude and place field location were randomly varied for each cell and day. Parameters are amplitude change probability (i.e., the probability that any given neuron would change its amplitude from one day to the next), magnitude of the amplitude change and remapping probability (i.e., probability for each neuron to change the location of its firing field). Right: Spatial tuning functions for simulated drift sorted for day 1. Simulated data shows time-dependent drift, similar to what is observed for real data. Edge angle dissimilarity for different levels of amplitude change probability. F=93.7-277.3, p=10^-4^-10^-5^, F=19.4-50.7, p=10^-20^-10^-33^ for condition and time*condition interaction effects vs. original data. **C.** Same as B but for the magnitude of the amplitude change. F=112.9-147.7, p=10^-4^-10^-5^, F=26.4-39.7, p=10^-2^4-10^-29^ for condition and time*condition interaction effects vs. original data. **D.** Same as B but for remapping probability. F=85.7-307.3, P=10^-4^-10^-5^, F=16.3-39.7, p=10^-18^-10^-29^ for condition and time*condition interaction effects vs. original data. **E.** Low edge angle dissimilarity persists when neurons with stable place fields (Pearson’s r >0.7) are removed from the analysis. F=164.2, p=10^-5^, F=37.1, t=10^-28^ for condition and time*condition interaction effects vs. control data with simulated drift (green) **F.** Stable edge angles persist when principal component analysis (PCA) is applied to the data first. F=679.2, p=10^-6^, F=15.1, t=10^-17^ for condition and time*condition interaction effects vs. control data with simulated drift (green). **G.** Same as D but for dPCA. F=1710, p=10^-7^, F=46.2, t=10^-31^ for condition and time*condition interaction effects vs. control data with simulated drift (green). All comparisons: 2-way repeated measures ANOVAs. Thick lines/shading: mean ± SEM, thin lines: individual mice (n=6).

**Supplemental Fig. 3:**
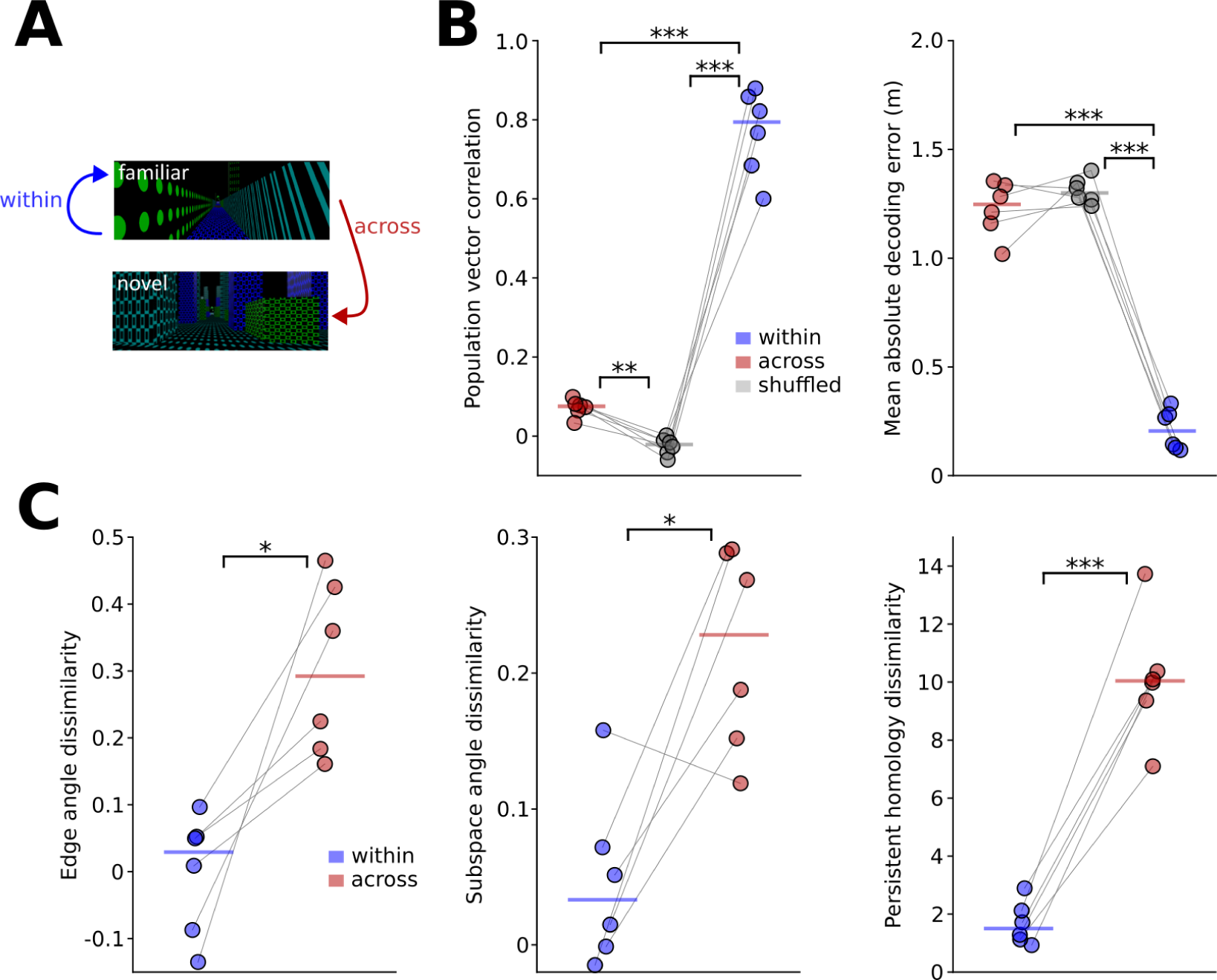
Non-preserved internal geometry across contexts. **A.** Schematic of the comparisons within the familiar environment (‘within’) and between the familiar and novel context (‘across’). **B.** Left: Population vector correlation is significantly lower across compared to within condition (t=-16.1, p=1.7*10^-5^, across vs. shuffled: t=6.7, p=0.003, within vs. shuffled: t=15.2, p=6.7*10^-5^, one-way repeated measures ANOVA followed by paired t-tests with Bonferroni correction), confirming remapping upon novel context exposure. Right: The error of a positional decoder is significantly larger when train and test data are obtained across environments compared to within the familiar environment (t=16.0, p=5.2*10^-5^, across vs. shuffled: t=-1.4, p=0.638, within vs. shuffled: t=-22.4, p=10^-6^, one-way repeated measures ANOVA followed by paired t-tests with Bonferroni correction). **C.** Dissimilarity metrics for edge angles (t=-4.0, p=0.011), subspace angles (t=-3.5, p=0.018) and persistent homology (t=-10.8, p=0.0001, paired t-tests) are lower for the within compared to the across condition, pointing to dissimilar internal geometries across environments.

**Supplemental Fig. 4:**
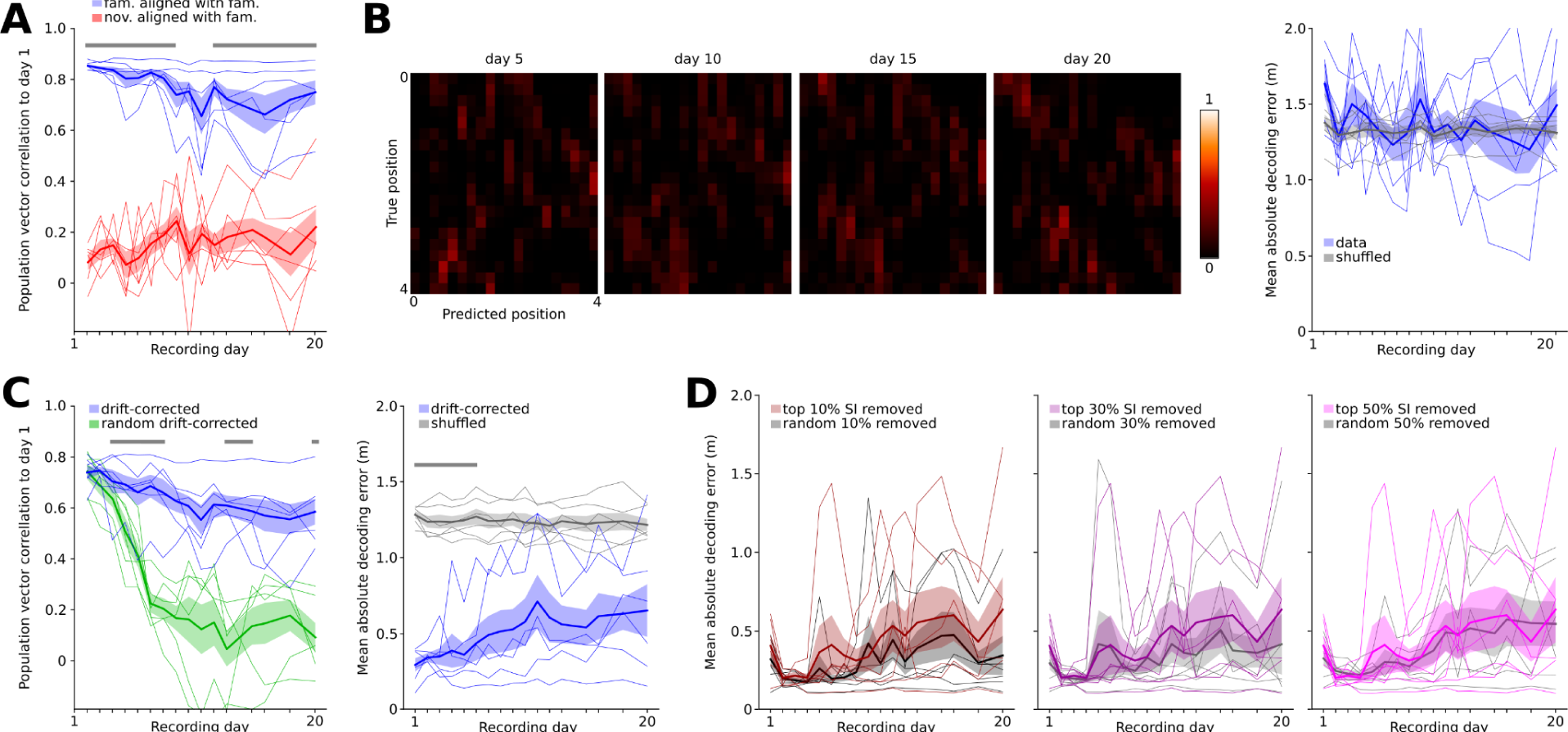
Downstream tuning correction across environments and as a function of spatial tuning. **A.** Population vector correlation in the novel environment aligned to the familiar arena compared to alignment within familiar. t=5.9-26.8, p=0.033-10^-5^. **B.** Predicting the animals’ position with decoders trained on the familiar and tested on the novel environment. Left: example confusion matrices. Right: predictions are indistinguishable from shuffled controls (F=0.14, p=0.724, effect of condition). **C.** Alignment analysis after first performing dimensionality reduction with PCA. Left: population vector correlation within the familiar environment compared to simulated data with random drift. t=5.5-14.3, p=0.045-0.0005. Right: positional predictions with decoders trained on day 1. t=6.1-15.1, p=0.028-0.0004. **D.** Unaltered positional predictions with decoders trained on day 1 after removing 10% (F=5.0, p=0.076), 30% (F=4.2, p=0.095) and 50% (F=1.1, p=0.348, effect of condition) of neurons with largest SI compared to control data sets with equal numbers of randomly removed neurons. 2-way repeated measures ANOVAs followed by paired *t* or Wilcoxon tests with Bonferroni correction. Grey bars: significant differences. Thick lines/shading: mean ± SEM, thin lines: individual mice (n=6).

**Supplemental Fig. 5:**
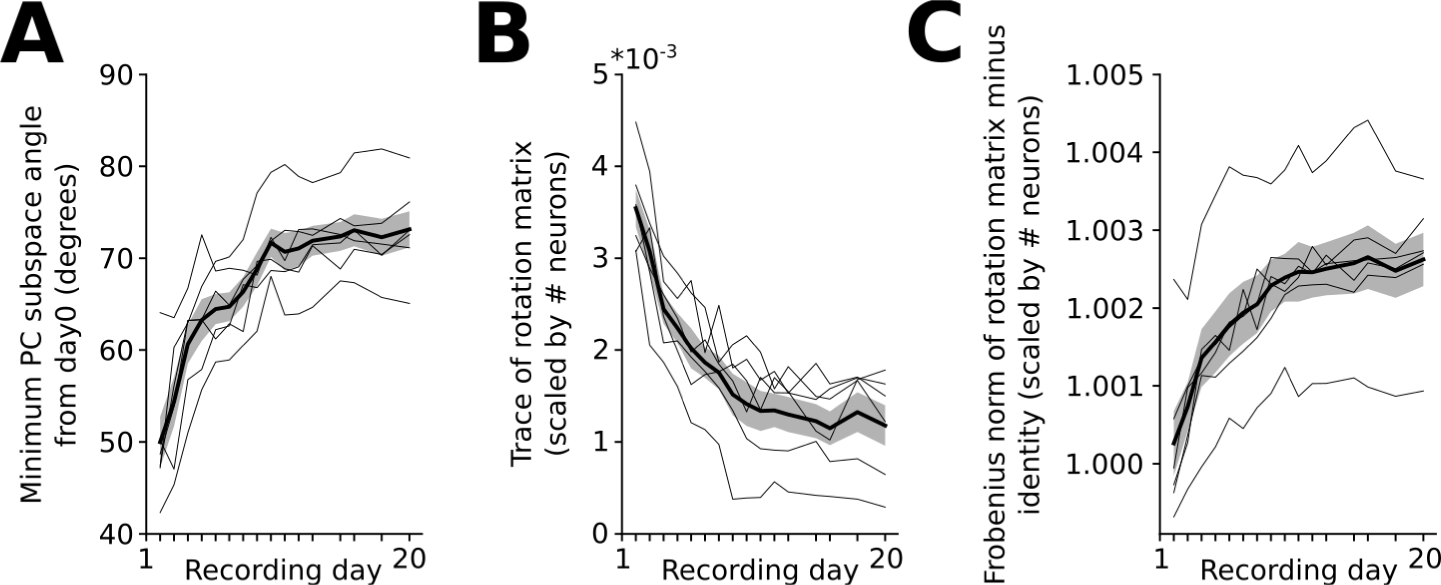
Alternative measures to assess the extent of rotation of the internal geometry across days. **A.** Minimum subspace angle (F=34.3, p=10^-27^). **B.** Trace of the alignment rotation matrix (F=51.3, p=10^-33^). **C.** Frobenius norm of the difference between the rotation matrix and the identity matrix (F=51.3, p=10^-33^). 1-way repeated measures ANOVAs. Thick lines/shading: mean ± SEM, thin lines: individual mice (n=6).

**Supplemental Fig 6:**
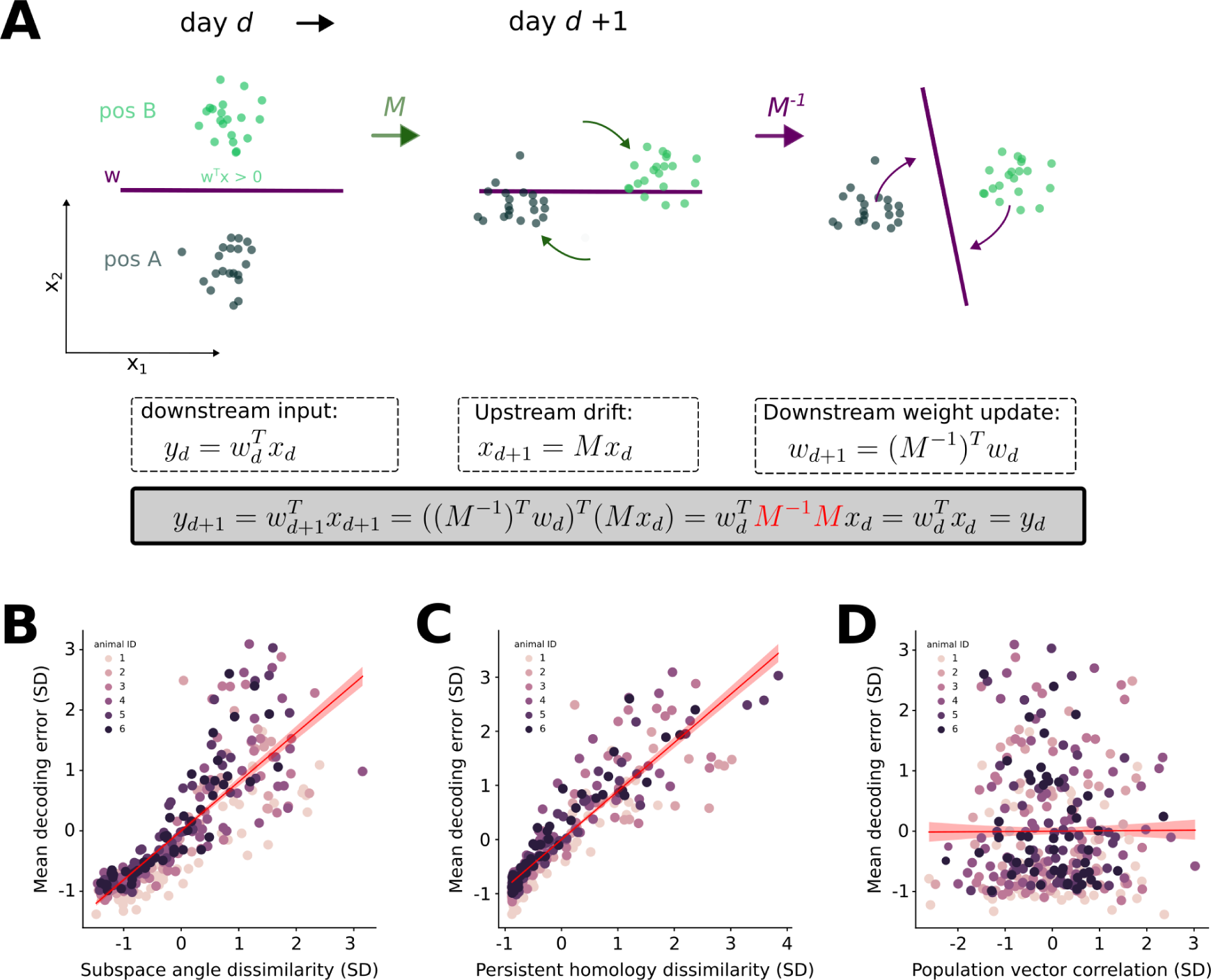
Downstream weight adaptations enable stable readout despite upstream representational drift. **A.** Schematic illustration using a toy example of two upstream neurons and one downstream neuron. Left: on day *d*, the activity of the upstream neurons (*x*_1_, *x*_2_) collectively encode two positions (pos A, dark green; pos B, light green). The downstream neuron with weights *w_d_*, activates if its inputs, *y_d_*, exceeds a threshold, which tends to be at position B. This creates a decision boundary (purple line) that separates the two positions. Middle: On a new day, the upstream representations drift in state space, modeled by applying a transformation matrix *M* to the upstream activity. Without downstream weight adaptation, the original decision boundary can no longer separate the positions. Right: The downstream neuron recovers the position selectivity by applying the inverse transformation to the weights, which rotates the decision boundary. **B.** Networks with larger subspace angle dissimilarity show larger decoding errors. β=0.857, p=10^-179^. **C.** Same as B but for persistent homology dissimilarity. β=0.908, p<10^-6^. **D.** Decoding errors do not correlate with the extent of drift quantified as population vector correlation. β=-0.068, p=0.236. B-D: linear mixed effect models. Thick lines/shading: mean ± SEM (n=6 mice).

## References

1. Robinson, N. T. M. et al. Targeted Activation of Hippocampal Place Cells Drives Memory-Guided Spatial Behavior. Cell 183, 1586–1599.e10 (2020).

2. Ziv, Y. et al. Long-term dynamics of CA1 hippocampal place codes. Nat. Neurosci. 16, 264–266 (2013).

3. Hainmueller, T. & Bartos, M. Parallel emergence of stable and dynamic memory engrams in the hippocampus. Nature 558, 292–296 (2018).

4. Geva, N., Deitch, D., Rubin, A. & Ziv, Y. Time and experience differentially affect distinct aspects of hippocampal representational drift. Neuron 0, (2023).

5. Gonzalez, W. G., Zhang, H., Harutyunyan, A. & Lois, C. Persistence of neuronal representations through time and damage in the hippocampus. Science 365, 821–825 (2019).

6. de Snoo, M. L., Miller, A. M., Ramsaran, A. I., Josselyn, S. A. & Frankland, P. W. Exercise accelerates place cell representational drift. Curr. Biol. CB 33, R96–R97 (2023).

7. Mankin, E. A. et al. Neuronal code for extended time in the hippocampus. Proc. Natl. Acad. Sci. 109, 19462–19467 (2012).

8. Dong, C., Madar, A. D. & Sheffield, M. E. J. Distinct place cell dynamics in CA1 and CA3 encode experience in new environments. Nat. Commun. 12, 2977 (2021).

9. Lee, J. S., Briguglio, J. J., Cohen, J. D., Romani, S. & Lee, A. K. The Statistical Structure of the Hippocampal Code for Space as a Function of Time, Context, and Value. Cell 183, 620–635.e22 (2020).

10. Schoonover, C. E., Ohashi, S. N., Axel, R. & Fink, A. J. P. Representational drift in primary olfactory cortex. Nature 594, 541–546 (2021).

11. Marks, T. D. & Goard, M. J. Stimulus-dependent representational drift in primary visual cortex. Nat. Commun. 12, 5169 (2021).

12. Dobler, Z. et al. Adapting and facilitating responses in mouse somatosensory cortex are dynamic and shaped by experience. Curr. Biol. 34, 3506–3521.e5 (2024).

13. Deitch, D., Rubin, A. & Ziv, Y. Representational drift in the mouse visual cortex. Curr. Biol. 31, 4327–4339.e6 (2021).

14. Chestek, C. A. et al. Single-Neuron Stability during Repeated Reaching in Macaque Premotor Cortex. J. Neurosci. 27, 10742–10750 (2007).

15. Driscoll, L. N., Pettit, N. L., Minderer, M., Chettih, S. N. & Harvey, C. D. Dynamic Reorganization of Neuronal Activity Patterns in Parietal Cortex. Cell 170, 986–999.e16 (2017).

16. Zhang, M. et al. The Representation of Decision Variables in Orbitofrontal Cortex is Longitudinally Stable. BioRxiv Prepr. Serv. Biol. 2024.02.16.580715 (2024) doi:10.1101/2024.02.16.580715.

17. Muysers, H. et al. A persistent prefrontal reference frame across time and task rules. Nat. Commun. 15, 2115 (2024).

18. Driscoll, L. N., Duncker, L. & Harvey, C. D. Representational drift: Emerging theories for continual learning and experimental future directions. Curr. Opin. Neurobiol. 76, 102609 (2022).

19. O’Keefe, J. & Dostrovsky, J. The hippocampus as a spatial map. Preliminary evidence from unit activity in the freely-moving rat. Brain Res. 34, 171–175 (1971).

20. Kentros, C. G., Agnihotri, N. T., Streater, S., Hawkins, R. D. & Kandel, E. R. Increased Attention to Spatial Context Increases Both Place Field Stability and Spatial Memory. Neuron 42, 283–295 (2004).

21. Zemla, R., Moore, J. J., Hopkins, M. D. & Basu, J. Task-selective place cells show behaviorally driven dynamics during learning and stability during memory recall. Cell Rep. 41, 111700 (2022).

22. Liberti, W. A., Schmid, T. A., Forli, A., Snyder, M. & Yartsev, M. M. A stable hippocampal code in freely flying bats. Nature 604, 98–103 (2022).

23. Zheng, Z. (Sam) et al. Perpetual step-like restructuring of hippocampal circuit dynamics. Cell Rep. 43, 114702 (2024).

24. Khatib, D. et al. Experience, not time, determines representational drift in the hippocampus. 2022.08.31.506041 Preprint at 10.1101/2022.08.31.506041 (2022).

25. Kaufman, M. T., Churchland, M. M., Ryu, S. I. & Shenoy, K. V. Cortical activity in the null space: permitting preparation without movement. Nat. Neurosci. 17, 440–448 (2014).

26. Khilkevich, A. et al. Brain-wide dynamics linking sensation to action during decision-making. Nature 634, 890–900 (2024).

27. Rule, M. E. et al. Stable task information from an unstable neural population. eLife 9, e51121 (2020).

28. Rule, M. E. & O’Leary, T. Self-healing codes: How stable neural populations can track continually reconfiguring neural representations. Proc. Natl. Acad. Sci. 119, e2106692119 (2022).

29. Sylte, O. C., Muysers, H., Chen, H.-L., Bartos, M. & Sauer, J.-F. Neuronal tuning to threat exposure remains stable in the mouse prefrontal cortex over multiple days. PLOS Biol. 22, e3002475 (2024).

30. Melbaum, S. et al. Conserved structures of neural activity in sensorimotor cortex of freely moving rats allow cross-subject decoding. Nat. Commun. 13, 7420 (2022).

31. Huang, L.-W., Torelli, F., Chen, H.-L. & Bartos, M. Context and space coding in mossy cell population activity. Cell Rep. 43, 114386 (2024).

32. Nieh, E. H. et al. Geometry of abstract learned knowledge in the hippocampus. Nature 595, 80–84 (2021).

33. Safaie, M. et al. Preserved neural dynamics across animals performing similar behaviour. Nature 1–7 (2023) doi:10.1038/s41586-023-06714-0.

34. Gallego, J. A., Perich, M. G., Chowdhury, R. H., Solla, S. A. & Miller, L. E. Long-term stability of cortical population dynamics underlying consistent behavior. Nat. Neurosci. 23, 260–270 (2020).

35. Kriegeskorte, N. & Diedrichsen, J. Peeling the Onion of Brain Representations. Annu. Rev. Neurosci. 42, 407–432 (2019).

36. Kriegeskorte, N. & Wei, X.-X. Neural tuning and representational geometry. Nat. Rev. Neurosci. 22, 703–718 (2021).

37. Harvey, S. E., Lipshutz, D. & Williams, A. H. What Representational Similarity Measures Imply about Decodable Information. Preprint at 10.48550/arXiv.2411.08197 (2024).

38. Fusi, S., Miller, E. K. & Rigotti, M. Why neurons mix: high dimensionality for higher cognition. Curr. Opin. Neurobiol. 37, 66–74 (2016).

39. Ostojic, S. & Fusi, S. Computational role of structure in neural activity and connectivity. Trends Cogn. Sci. 28, 677–690 (2024).

40. Rubin, A. et al. Revealing neural correlates of behavior without behavioral measurements. Nat. Commun. 10, 4745 (2019).

41. Lee, J. Q., Keinath, A. T., Cianfarano, E. & Brandon, M. P. Identifying representational structure in CA1 to benchmark theoretical models of cognitive mapping. Neuron 113, 307–320.e5 (2025).

42. Bernardi, S. et al. The Geometry of Abstraction in the Hippocampus and Prefrontal Cortex. Cell 183, 954–967.e21 (2020).

43. Courellis, H. S. et al. Abstract representations emerge in human hippocampal neurons during inference. Nature 1–9 (2024) doi:10.1038/s41586-024-07799-x.

44. Qin, S. et al. Coordinated drift of receptive fields in Hebbian/anti-Hebbian network models during noisy representation learning. Nat. Neurosci. 26, 339–349 (2023).

45. Remme, M. W. H. et al. Hebbian plasticity in parallel synaptic pathways: A circuit mechanism for systems memory consolidation. PLOS Comput. Biol. 17, e1009681 (2021).

46. Rais, C., Maheu, M. & Wiegert, J. S. Schaffer commissural inputs regulate the stability of CA1 spatial representation. 2025.01.24.634689 Preprint at 10.1101/2025.01.24.634689 (2025).

47. Basu, J. & Siegelbaum, S. A. The Corticohippocampal Circuit, Synaptic Plasticity, and Memory. Cold Spring Harb. Perspect. Biol. 7, a021733 (2015).

48. Savelli, F. & Knierim, J. J. Hebbian Analysis of the Transformation of Medial Entorhinal Grid-Cell Inputs to Hippocampal Place Fields. J. Neurophysiol. 103, 3167–3183 (2010).

49. Solstad, T., Moser, E. I. & Einevoll, G. T. From grid cells to place cells: A mathematical model. Hippocampus 16, 1026–1031 (2006).

50. Udakis, M., Pedrosa, V., Chamberlain, S. E. L., Clopath, C. & Mellor, J. R. Interneuron-specific plasticity at parvalbumin and somatostatin inhibitory synapses onto CA1 pyramidal neurons shapes hippocampal output. Nat. Commun. 11, 4395 (2020).

51. Attardo, A., Fitzgerald, J. E. & Schnitzer, M. J. Impermanence of dendritic spines in live adult CA1 hippocampus. Nature 523, 592–596 (2015).

52. Barack, D. L. & Krakauer, J. W. Two views on the cognitive brain. Nat. Rev. Neurosci. 22, 359–371 (2021).

53. Ebitz, R. B. & Hayden, B. Y. The population doctrine in cognitive neuroscience. Neuron 109, 3055–3068 (2021).

54. Mante, V., Sussillo, D., Shenoy, K. V. & Newsome, W. T. Context-dependent computation by recurrent dynamics in prefrontal cortex. Nature 503, 78–84 (2013).

55. Wang, J., Narain, D., Hosseini, E. A. & Jazayeri, M. Flexible timing by temporal scaling of cortical responses. Nat. Neurosci. 21, 102–110 (2018).

56. Pachitariu, M. et al. Suite2p: beyond 10,000 neurons with standard two-photon microscopy. 061507 Preprint at 10.1101/061507 (2017).

57. Dombeck, D. A., Harvey, C. D., Tian, L., Looger, L. L. & Tank, D. W. Functional imaging of hippocampal place cells at cellular resolution during virtual navigation. Nat. Neurosci. 13, 1433–1440 (2010).

58. Skaggs, W., McNaughton, B. & Gothard, K. An Information-Theoretic Approach to Deciphering the Hippocampal Code. in Advances in Neural Information Processing Systems vol. 5 (Morgan-Kaufmann, 1992).

59. Kobak, D. et al. Demixed principal component analysis of neural population data. eLife 5, e10989 (2016).

60. Tralie, C., Saul, N. & Bar-On, R. Ripser.py: A Lean Persistent Homology Library for Python. J. Open Source Softw. 3, 925 (2018).

61. Rybakken, E., Baas, N. & Dunn, B. Decoding of Neural Data Using Cohomological Feature Extraction. Neural Comput. 31, 68–93 (2019).

62. Chaudhuri, R., Gerçek, B., Pandey, B., Peyrache, A. & Fiete, I. The intrinsic attractor manifold and population dynamics of a canonical cognitive circuit across waking and sleep. Nat. Neurosci. 22, 1512–1520 (2019).

63. Giusti, C., Pastalkova, E., Curto, C. & Itskov, V. Clique topology reveals intrinsic geometric structure in neural correlations. Proc. Natl. Acad. Sci. 112, 13455–13460 (2015).

64. Dabaghian, Y., Mémoli, F., Frank, L. & Carlsson, G. A Topological Paradigm for Hippocampal Spatial Map Formation Using Persistent Homology. PLOS Comput. Biol. 8, e1002581 (2012).

65. Gardner, R. J. et al. Toroidal topology of population activity in grid cells. Nature 602, 123–128 (2022).

66. Hermansen, E., Klindt, D. A. & Dunn, B. A. Uncovering 2-D toroidal representations in grid cell ensemble activity during 1-D behavior. Nat. Commun. 15, 5429 (2024).

67. Kawahara, D. & Fujisawa, S. Advantages of Persistent Cohomology in Estimating Animal Location From Grid Cell Population Activity. Neural Comput. 36, 385–411 (2024).

